# MUTE drives asymmetric divisions to form stomatal subsidiary cells in Crassulaceae succulents

**DOI:** 10.1101/2024.12.27.630159

**Authors:** Xin Cheng, Heike Lindner, Lidia Hoffmann, Antonio Aristides Pereira Gomes Filho, Paola Ruiz Duarte, Susanna F Boxall, Yigit Berkay Gündogmus, Jessica H Pritchard, Sam Haldenby, Matthew Gemmell, Alistair Darby, Miro Läderach, James Hartwell, Michael T Raissig

## Abstract

Amongst the evolutionary innovations of many succulents is a photosynthetic lifestyle, where stomatal gas exchange is decoupled from light-dependent carbon fixation. Stomatal complexes in the emerging succulent model *Kalanchoë laxiflora* consist of two guard cells surrounded by three anisocytic subsidiary cells (SCs). Here, we show that these SCs shuttle ions and thus likely support stomatal movements. Furthermore, gene editing, reporter lines and protein overexpression implicate the stomatal transcription factor MUTE in facilitating additional rounds of asymmetric divisions that form SCs in succulents. This is opposite to the role of MUTE in *Arabidopsis thaliana*, where it stops rather than induces asymmetric divisions, but reminiscent of MUTE’s SC-related function in grasses. Together, our work deciphers an intricate genetic mechanism that generates innovative stomatal morphology in Crassulaceae succulents.

## Introduction

Land plants evolved numerous anatomical and physiological features to thrive under harsh terrestrial conditions. Amongst them are sealed leaves preventing desiccation that are interspersed with adjustable breathing pores (=stomata) that enable plant-atmosphere gas exchange (Hetherington and Woodward, 2003). Additional evolutionary innovations like plant organ succulence and Crassulacean Acid Metabolism (CAM), a highly water-use efficient photosynthetic life-style, further unlocked extremely water-scarce environments (Griffiths and Males, 2017; Heyduk, 2022; Sage et al., 2023). CAM plants open stomata primarily at night and fix carbon dioxide (CO_2_) into an organic acid intermediate stored in vacuoles. During the day, stomata are mostly closed, the storage intermediate is decarboxylated, and the free CO_2_ is assimilated in the Calvin-Benson-Bassham cycle (Borland et al., 2009; Osmond, 1978). Most succulent plants employ CAM photosynthesis and many CAM plants form stomata, where the central guard cells (GCs) are flanked or surrounded by putative subsidiary cells (SCs; Males and Griffiths, 2017). Stomata in the emerging leaf succulent CAM model *Kalanchoë laxiflora* (Boxall et al., 2020; Hartwell et al., 2016), for example, consist of two kidney-shaped GCs surrounded by three unequally sized (=anisocytic), circularly arranged SCs (Fig. 1A and B). These SCs are distinct in size and shape compared to the puzzle-shaped pavement cells (PCs), and thus fulfil the original definition of stomatal SCs (Esau, 1965). Yet, even though SCs are overrepresented in CAM plants, it remained unclear if they indeed serve as helper cells to support stomatal opening and closing like in grasses (Durney et al., 2023; Franks and Farquhar, 2007; Raissig et al., 2017).

**Figure 1.**
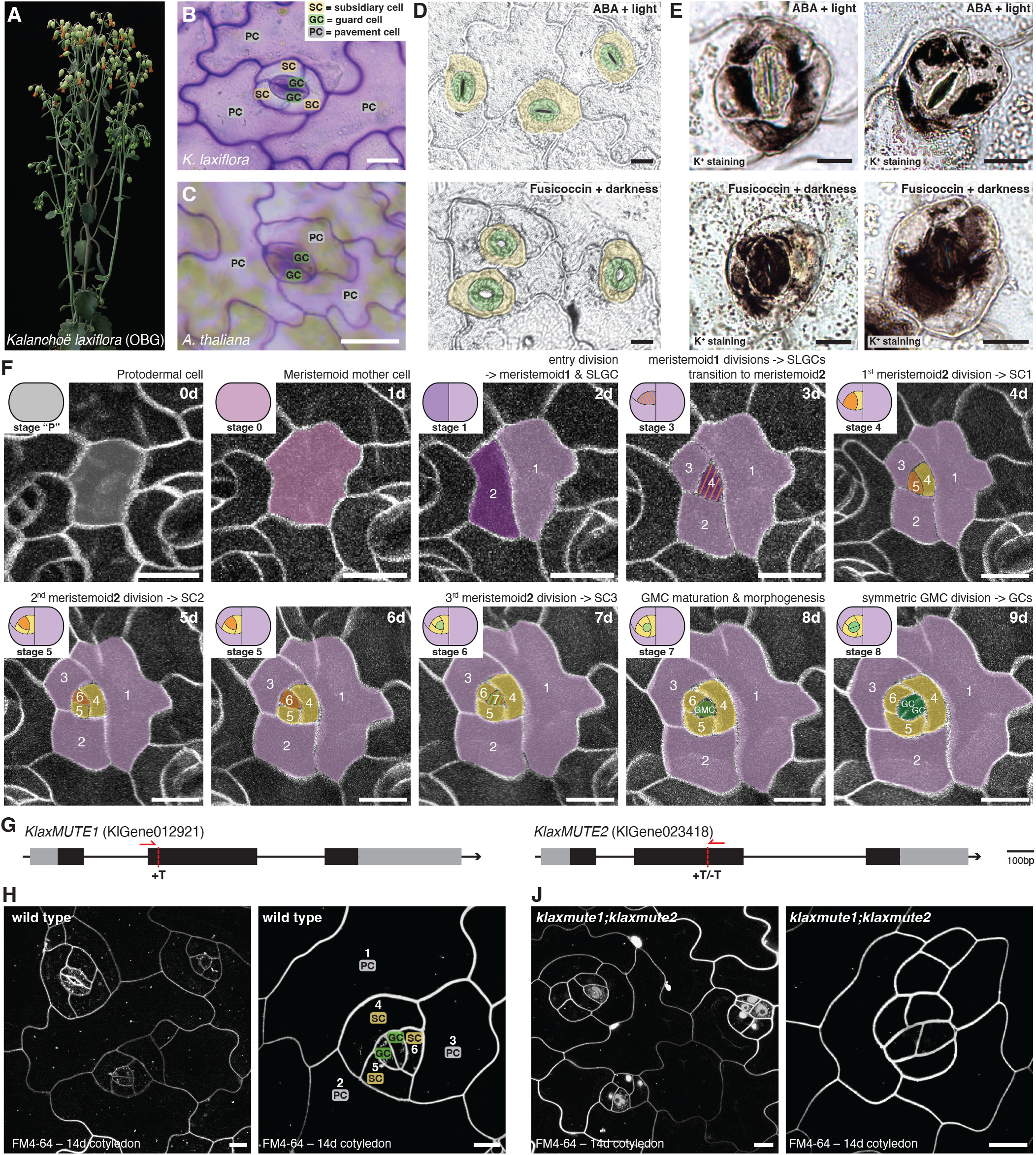
*MUTE*-dependent amplifying divisions form functionally relevant subsidiary cells in *Kalanchoë laxiflora*. (A) Flowering *K. laxiflora*, an emerging succulent model plant. (B) *K. laxiflora* stomata consisting of two central guard cells (GCs) surrounded by three anisocytic subsidiary cells (SCs). (C) Two-celled stomata of *A. thaliana* without SCs. (D) Abscisic acid (ABA, top) and Fusicoccin treatment (bottom) close and open stomata, respectively. GCs are false-colored in green and SCs in yellow. (E) Potassium (K^+^)-staining shows K^+^ in SCs when stomata are closed and in GCs when stomata are open. (F) 10-day time-lapse imaging of plasma membrane marker line (*35Sp:mCherry-AtPIP1;4*) reveals distinct phases of asymmetric divisions before GCs are formed. Stages and days are indicated. Cell types are false-colored; stomatal lineage ground cells (SLGCs) are in lilac, SCs in yellow, guard mother cells (GMCs) and GCs in green, meristemoid1 in purple, meristemoid2 in orange. (G) Gene models of the two duplicated *KlaxMUTE* homologs; gene-editing sites and guide RNAs are indicated. (H, J) Developing 14d-old cotyledons in wild type (H) and *klaxmute1;klaxmute2* double mutants (J). Cell membranes visualized using FM4-64. Scale bars, 20 µm.

Unlike many CAM plants, the C_3_ eudicot and Brassi-caceae model *Arabidopsis thaliana* does not form SCs (Fig. 1C). Consequently, the two GCs are surrounded by puzzle-shaped PCs only (Fig. 1C). In *A. thaliana*, three basic Helix-Loop-Helix (bHLH) transcription factors guide stomatal development. *AtSPEECHLESS* (*AtSPCH*) determines stomatal precursors (=meristemoids) and drives asymmetric divisions (MacAlister et al., 2007). *AtMUTE* ends asymmetric divisions, establishes the guard mother cell (GMC) and induces the symmetric division forming the GC pair (Han et al., 2022, 2018; Pillitteri et al., 2007). Finally, *AtFAMA* represses symmetric division potential and controls GC differentiation (Ohashi-Ito and Bergmann, 2006). While this bHLH module is conserved from mosses to flowering plants (Chater et al., 2017), it was amplified, sub-functionalized, modified and rewired to generate different stomatal morphologies (Chater et al., 2016; Raissig et al., 2016). In grasses, for example, MUTE is not required for GMC identity but rather acquired cell-to-cell mobility to establish subsidiary mother cells (SMCs) in cells adjacent to the GMC, which then divide asymmetrically to contribute perigene SCs (i.e. from a distinct cell lineage to those that lead to GCs; Raissig et al., 2017; Wang et al., 2019; Wu et al., 2019). In *K. laxiflora*, though, both SCs and GCs are formed by the same precursor cell (= meristemoid), which seems to undergo many rounds of spiralling amplifying divisions, resulting in mesogene SCs (i.e. from the same lineage as GCs; Cheng and Raissig, 2023; Rudall et al., 2013). Yet how often, and in which pattern, meristemoids divide, why some divisions yield SCs and which genetic programs control these divisions remained elusive.

Here, we use the eudicot succulent CAM model *K. laxiflora* (Oxford Botanical Garden, OBG accession) to determine the functional relevance of SCs, the division patterns that form SCs and the genetic program that enables SC divisions. We sequenced *K. laxiflora* OBG’s diploid genome, established efficient transgenesis protocols, and optimized horticultural protocols to efficiently induce flowering and viable seed production and reduce the life cycle to less than 6 months. Together, this makes *K. laxiflora* OBG an ideal model system to dissect developmental mechanisms in a leaf succulent plant. Potassium (K^+^) staining revealed K^+^ shuttling between GCs and SCs during stomatal movements, suggesting that the SCs are indeed physiological helper cells, as in grasses (Raschke and Fellows, 1971). Time-lapse imaging of developing leaves showed that additional rounds of amplifying divisions, which do not occur in *A. thaliana*, generated the mesogenous, anisocytic SCs. Mutant, reporter and overexpression analysis of the duplicated *KlaxMUTE1* and *KlaxMUTE2* genes provided strong support for the conclusion that *Kalanchoë* MUTEs control an asymmetric division program that generates SCs. *KlaxMUTEs*’ novel function in driving SC divisions was species-context-dependent and could not be transferred to *A. thaliana*. Finally, transcriptome analysis indicated that *KlaxMUTE1* activated specific cell-cycle programs associated with asymmetric rather than symmetric cell divisions and delayed GC differentiation. Together, our work uncovered, for the first time, the intricate genetic mechanisms that govern the formation of functionally relevant, anisocytic SCs in the Crassulaceae family that likely play a pivotal role in the CAM photosynthetic lifestyle.

### K^+^ shuttling during stomatal movements suggests subsidiary cells to be functionally relevant

To determine if *Kalanchoë* SCs are indeed helper cells that contribute to stomatal opening and closing, we precipitated the major stomatal osmolyte potassium (K^+^) in open and closed stomata in *K. laxiflora* (Fig. 1D and E). Incubation of epidermal leaf peels with Fusicoccin and ABA successfully opened and closed stomata, respectively (Fig. 1D). Upon treatments, K^+^ precipitation showed that K^+^ primarily resides in SCs in closed stomata (Fig. 1E) and moves to GCs in open stomata (Fig. 1E). This strongly suggests that, much like in grasses, SCs contribute to osmolyte shuttling and thus turgor control of stomatal cells (Raschke and Fellows, 1971). Quantification of K^+^ levels in open and closed stomata, however, suggested that residual K^+^ remains in SCs even when stomata are open (Fig. S1).

### Additional rounds of asymmetric meristemoid divisions generate subsidiary cells

Mature *Kalanchoë* stomata consist of two paired GCs surrounding the pore and three unequally sized and circularly arranged SCs. In addition, the complete stomatal complex is surrounded by 2 to 3 PCs (Fig. 1B). We previously suggested that protodermal cells undergo 5 to 6 rounds of stem-cell-like asymmetric cell divisions (ACDs) following a Fibonacci spiral pattern, where the first 2 to 3 divisions generate stomatal lineage ground cells (SLGCs) and the following three ACDs generate SCs (Cheng and Raissig, 2023). To unequivocally describe stomatal development in *K. laxiflora*, we employed manual time-lapse imaging of developing leaves expressing a plasma membrane reporter gene (*35Sp:mCherry-AtPIP1;4*) to follow developing stomatal complexes over 10 days (Fig. 1F, Movie S1). Indeed, a meristemoid mother cell executed an initial entry division (stage 1; Fig. 1F). The smaller cell then divided asymmetrically two more times forming SLGCs while maintaining a small, stem-cell-like meristemoid (stage 2 and 3, Fig. 1F). Much like in *A. thaliana*, SLGCs can directly differentiate into PCs or execute a spacing division to establish an additional meristemoid (Fig. S2; Movie S2). After the entry division and two SLGC divisions, the meristemoid divided three additional times to produce SCs, which could not execute further spacing divisions (stage 4 to 6, Fig. 1F). After three SC divisions, the meristemoid then adopted GMC identity, rounded up (stage 7), and divided symmetrically to form GCs (stage 8, Fig. 1F). We therefore suggest two distinct meristemoid stages; a meristemoid1 stage that produces SLGCs and a meristemoid2 stage that produces SCs (Fig. 1F). However, the developmental mechanisms and genetic modules that control the meristemoid1-to-meristemoid2 transition and the SLGC versus SC divisions remained unknown.

### *klaxmute1;klaxmute2* cannot form functional stomatal complexes and shows aberrant asymmetric divisions

In *A. thaliana*, the *MUTE* transcription factor ends meristemoid ACDs and controls the meristemoid-to-GMC transition (Han et al., 2022, 2018; Pillitteri et al., 2007). In grasses, however, *MUTE* controls SC formation and controls GMC division orientation (Raissig et al., 2017; Spiegelhalder et al., 2024; Wang et al., 2019). In *K. laxiflora*, the duplicated *MUTE* homologs *KlaxMUTE1* and *KlaxMUTE2* show an extended C-terminal amino acid tail as is the case for all grass *MUTE* homologs, but unlike most other eudicot homologs, suggesting a function more similar to grasses rather than *A. thaliana* (Fig. S3). We used gene editing to induce mutations in both *KlaxMUTE* homologs (Fig. 1G). While the single mutants did not show a mutant phenotype (Fig. S4A), *klaxmute1;klaxmute2* double mutants arrested growth as young seedlings and failed to form mature stomata (Fig. 1H and J, Fig. S4B). Instead, we observed additional, misoriented ACDs that violated the Fibonacci spiral pattern (Fig. 1H and J). This indicated a role for the *KlaxMUTEs* beyond controlling the meristemoid-to-GMC transition observed in *A. thaliana*.

### *KlaxMUTE1* and *KlaxMUTE2* reporter genes are expressed during subsidiary cell divisions

To observe when and where *KlaxMUTE1* and *KlaxMUTE2* are expressed, we generated transcriptional and translational reporter lines for both *KlaxMUTE* homologs. Strikingly, both transcriptional reporters (*KlaxMUTE1p:m-Cit-eGFP*^*nls*^ and *KlaxMUTE2p:mCit-eGFP*^*nls*^) started to be expressed in meristemoids after the first three ACDs (i.e. entry division and two SLGC divisions) concomitant with the meristemoid1-to-meristemoid2 transition (stage 3; Fig. 2A to C). The two *KlaxMUTE* promoters remained active throughout all meristemoid2 divisions that generate SCs and peaked in GMCs (stage 4 to 6-7; Fig. 2B and C). A similar expression window was observed with translational reporters (*KlaxMUTE1p:mCit-KlaxMUTE1, KlaxMUTE2p:m-Cit-KlaxMUTE2*, Fig. 2D and E). Weak expression of both translational reporter lines was first detected during the meristemoid1-to-meristemoid2 transition (stage 3) and was maintained until after the symmetric GMC division (Fig. 2D and E). Both translational reporters increased in expression strength during the SC divisions, yet seemed to be downregulated post-mitotically in the newly formed SCs–potentially to prevent ectopic divisions in SCs. During GMC formation (stage 6-7), we observed strong nuclear and weak cytoplasmic signals in GMCs and a re-appearing signal in SCs. The signal intensity decreased quickly in all cells after the symmetric GMC division (stage 8, Fig. 2D and E), but remained longer in divided GCs and sometimes weak signal remained in the smallest SC (Fig. S5). Together, the expression window of *KlaxMUTEs* coincided exactly with all SC divisions, suggesting a novel role that facilitates the formation of anisocytic SCs.

**Figure 2.**
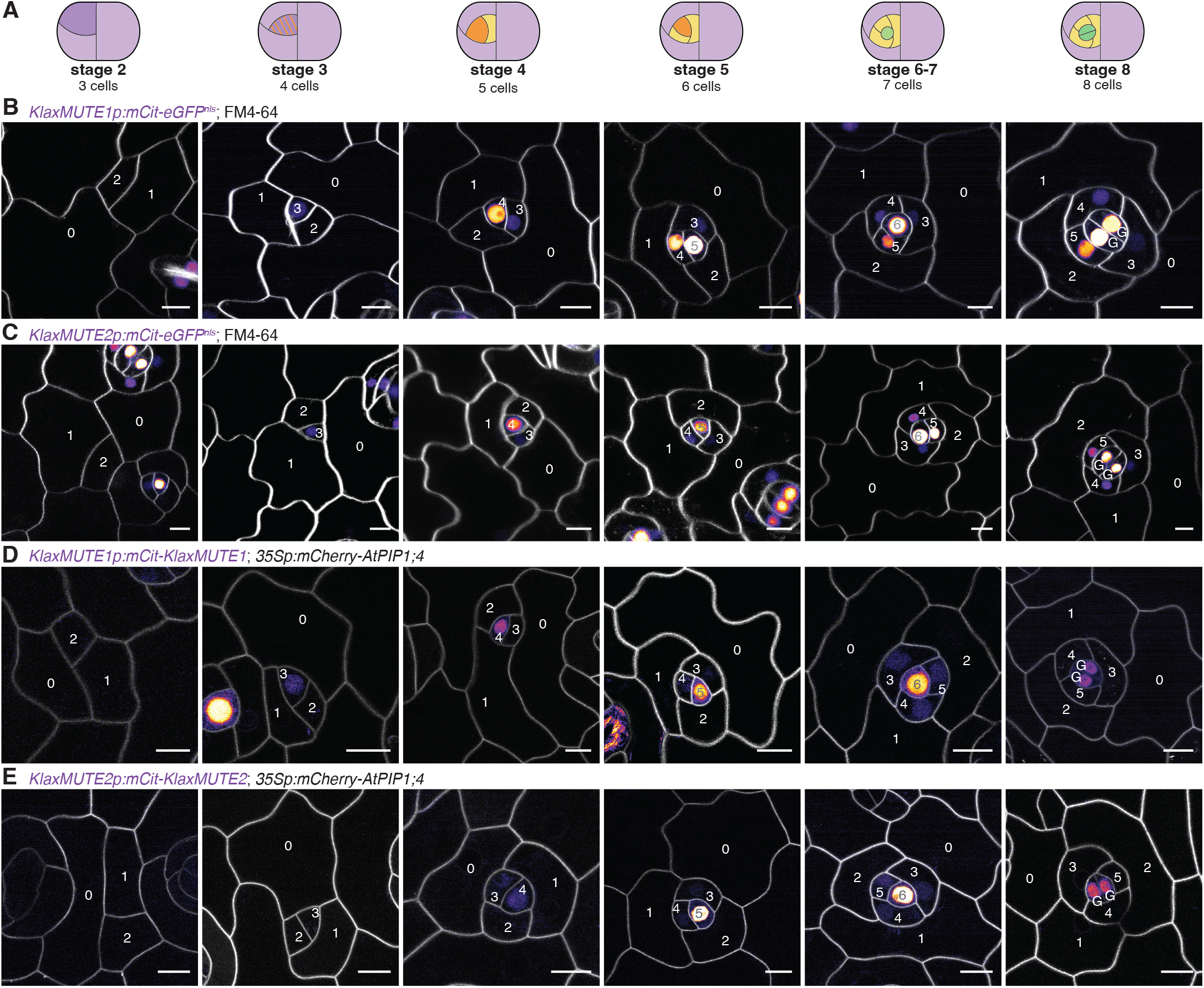
*KlaxMUTE1* and *KlaxMUTE2* are expressed throughout subsidiary cell divisions. (A) Different developmental stages of stomata (stage 2 until stage 8) shown below. (B, C) Confocal microscopy images of transcriptional reporters *KlaxMUTE1p:mCi-trine-eGFP*^*nls*^ (B) and *KlaxMUTE2p:mCitrine-eGFP*^*nls*^ (C). Cell membranes were stained with FM4-64. (D, E) Confocal microscopy images of translational reporters *KlaxMUTE1p:mCitrine-KlaxMUTE1* (D) and *KlaxMUTE2p:mCitrine-KlaxMUTE2* (E) also expressing the plasma membrane marker *35Sp:mCherry-AtPIP1;4*. Scale bars, 10 µm.

### Overexpression of *KlaxMUTEs* induces ectopic asymmetric divisions

The key experiment in *A. thaliana* that firmly determined MUTE’s role in ending ACDs and committing GMC identity was its overexpression. Overexpressing *AtMUTE* produced a leaf epidermis solely consisting of paired GCs (Pillitteri et al., 2007). However, overexpressing *KlaxMUTE1* (*35Sp:mCit-KlaxMUTE1 = KlaxMUTE1*^*OE*^) or *KlaxMUTE2* (*35Sp:mCit-KlaxMUTE2 = KlaxMUTE2*^*OE*^) in *K. laxiflora* caused a severe overabundance of small cells in the mature leaf epidermis and a striking absence of puzzle-shaped PCs (Fig. 3A to C). The total cell density was increased fourfold and three-fold in *KlaxMUTE1*^*OE*^ and *KlaxMUTE2*^*OE*^, respectively (Fig. 3D). The stomatal density, on the other hand, was increased in *KlaxMUTE1*^*OE*^ only, and slightly and non-significantly decreased in *KlaxMUTE2*^*OE*^ (Fig. 3E). Furthermore, more than 35% and 20% of stomatal complexes formed more than 3 SCs in *KlaxMUTE1*^*OE*^ and *KlaxMUTE2*^*OE*^, respectively (Fig. 3F). Together, this strongly suggested that the *KlaxMUTEs* induce an asymmetric division program to form SCs rather than establishing GMC identity and ending ACDs, as is the case in *A. thaliana*. Indeed, observing young developing leaves (0.5 cm) showed numerous ectopic ACDs and many more, and much smaller, cells in the leaf epidermis compared to similar-sized wild-type leaves (Fig. S6A to C). Despite these massive changes to leaf epidermal development, the overexpression lines grew reasonably well but formed slightly cupshaped, curved leaves, which might be caused by imbalanced abaxial and adaxial tissue growth patterns as a consequence of ectopic divisions (Fig. S6D).

**Figure 3.**
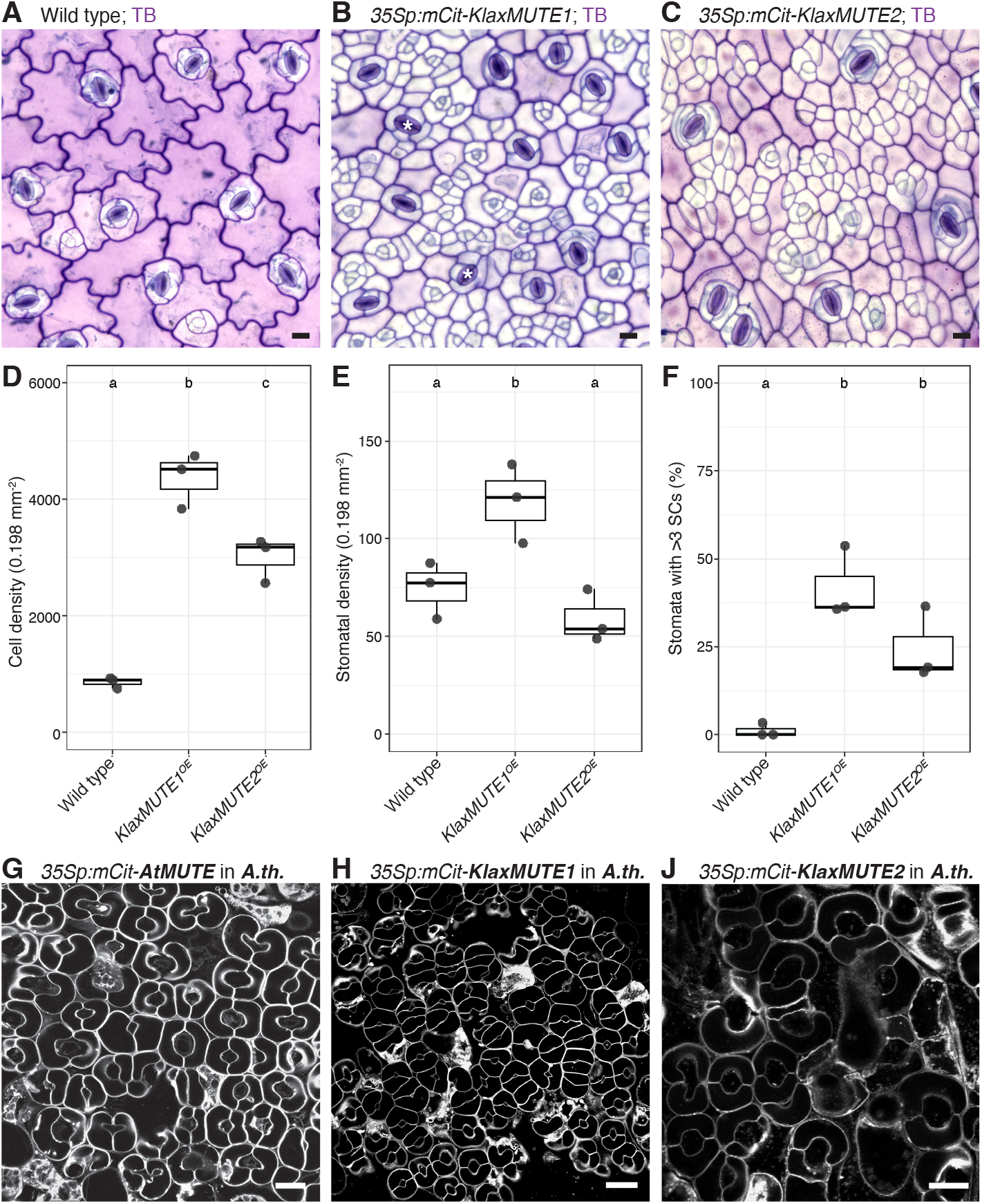
Overexpression of *KlaxMUTE1* and *KlaxMUTE2* induces ectopic asymmetric divisions in *K. laxiflora*. (A – C) Brightfield images of Toluidine Blue-stained epidermal peels of mature *K. laxiflora* leaves in wild type (A), *35Sp:mCitrine-KlaxMUTE1* (B) and *35S:mCitrine-KlaxMUTE2* (C). Scale bar, 20 µm. (D – F) Quantification of *Klax-MUTE1* and *KlaxMUTE2* overexpression phenotypes; cell density (D), stomatal density (E), and fraction of stomatal complexes with more than three subsidiary cells (SCs, F). Counted were three individuals per genotype, 1500-7780 total cells and 100-212 stomata per genotype. One-way ANOVA followed by Tukey post-hoc test, letters indicate significance groups. (G-J) Confocal images of FM4-64-stained cotyledons of *MUTE* overexpression lines in *A. thali-ana*; *35Sp:mCitrine-AtMUTE* (G), *35Sp:m-Citrine-KlaxMUTE1* (H), and *35Sp:mCi-trine-KlaxMUTE2*(J). Scale bar, 20 µm.

While we cannot fully determine the identity of the ectopically formed small cells, circumstantial evidence suggested them to be SC-like cells. First, in Toluidine blue-stained epidermal peels of mature wild-type leaves, the three epidermal cell types have very specific staining patterns; GCs were purple, SCs were white (i.e. unstained), and PCs were pink (Fig. 3A). The numerous ectopic small cells in the overexpression lines mostly remained white indicating SC-like characteristics (Fig. 3B and C). Furthermore, PCs are known to endoreduplicate and are thus of higher-order ploidies (Melaragno et al., 1993). Ploidy analysis of epidermal nuclei (i.e. of isolated peels) indicated fewer polyploid nuclei in the overexpression lines compared to wild type, further suggesting a smaller fraction of PCs and a larger fraction of SC-like cells (Fig. S7).

Together, we conclude that Crassulaceae *MUTEs* control the transition from the SLGC-producing meristemoid1 to the SC-producing meristemoid2 and promote additional rounds of amplifying divisions required to form three anisocytic SCs.

### The novel role of *KlaxMUTEs* is species context-dependent

Next, we wanted to test if the ACD-promoting functionality of *K. laxiflora* MUTE was intrinsic to the protein, and could thus be transferred to *A. thaliana*. However, overexpression of either *AtMUTE* (*35Sp:mCit-AtMUTE*), *KlaxMUTE1* (*35Sp:mCit-KlaxMUTE1*) or *KlaxMUTE2* (*35Sp:mCit-KlaxMUTE2*) in *A. thaliana* solely induced paired GC formation and not ectopic ACDs (Fig. 3G to J). Similarly, overexpressing *AtMUTE* in *K. laxiflora* induced additional ACDs rather than paired GC formation like in *A. thaliana* (Fig. S6E). Together, this suggested that either species-specific heterodimerization partners, or a diversified genetic program downstream of *MUTE*, underlie the divergent function of *MUTE* in *K. laxiflora*.

### *KlaxMUTE1* induces meristemoid2-specific cell division and identity programs

To determine potential downstream programs activated by *KlaxMUTE1*, we analyzed the transcriptome of mature (7^th^) leaves of *KlaxMUTE1*^*OE*^ (Fig. 4A). Compared to mature wild-type leaves, thousands of genes were differentially expressed (2759 up and 1762 down, log2FoldChange >2, padj < 0.1; Fig. 4A, Table S3). The most upregulated gene was the overexpressed *KlaxMUTE1* itself, which also upregulated its sister gene *KlaxMUTE2* (Fig. 4A, Table S3). We then identified homologs of key stomatal regulators described in *A. thaliana* and determined if some of these were differentially regulated by overexpressing *KlaxMUTE1* compared to induced overexpression of *AtMUTE* in *A. thaliana* (*iMUTE*) using a dataset published previously (Fig. 4B, Table S3; Han et al., 2018). Strikingly, core developmental factors associated with GC differentiation like *KlaxFAMA2* and *KlaxFOUR-LIPS (KlaxFLP)* were downregulated by *KlaxMUTE1*^*OE*^, which was contrary to *A. thaliana*, where *AtFAMA* and *AtFLP*, and thus GC differentiation, was promoted by *AtMUTE* (Fig. 4B). In addition, two CycD3 homologs of the significantly amplified CycD3 gene family (9 genes in *K. laxiflora*, 3 genes in *A. thaliana*), which are associated with ACDs (Dewitte et al., 2007), were strongly induced in *KlaxMUTE1*^*OE*^ (Fig. 4B). In *A. thaliana*, CycD3 genes are bound and activated by *AtSPCH* instead, and are post-translationally inhibited in an AtMUTE-dependent manner during the meristemoid-to-GMC transition (Adrian et al., 2015; Dewitte et al., 2007; Han et al., 2022, 2018; Lau et al., 2014). Also the amplified CycD5 family showed two induced and one downregulated gene further indicating that *KlaxMUTE1* induces a meristemoid2-specific cell division program. Other stomatal regulators behaved similarly in response to *MUTE* overexpression in both *K. laxiflora* and *A*.

**Figure 4.**
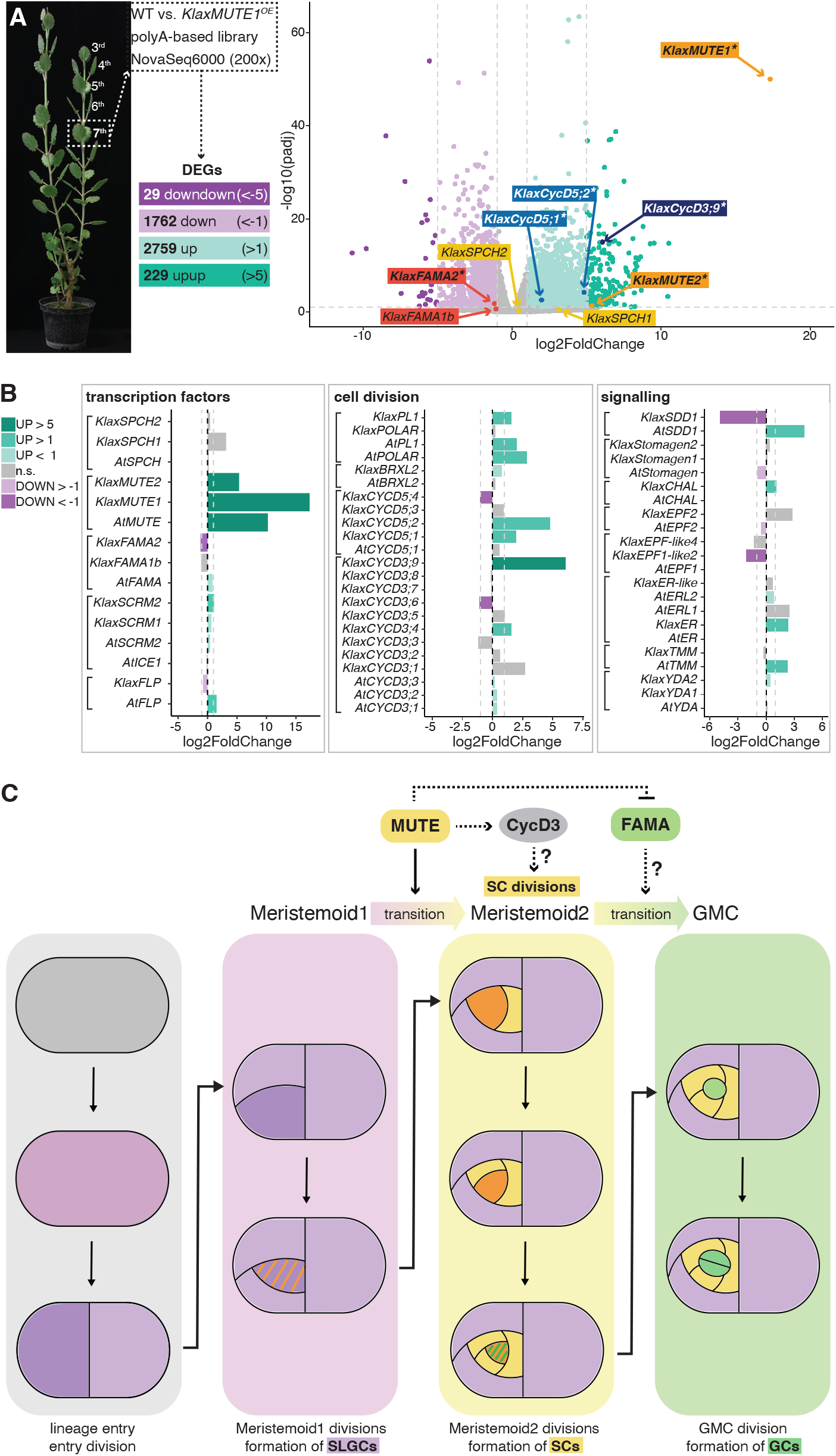
RNA-sequencing of *KlaxMUTE1* overexpression line. (A) Sampled leaf pairs and volcano plot of upregulated and down-regulated genes. Number of differentially expressed genes (DEGs) and selected candidate genes are indicated (significant DEGs are printed in bold and with an asterisk). (B) Comparison of upregulation and downregulation of stomatal developmental genes by induced *AtMUTE* overexpression in *A. thaliana* (*iMUTE*; data from Han et al., 2018) and *KlaxMUTE1* overexpression in *K. laxiflora*; stomatal transcription factors on the left, cell cycle-related genes in the middle, signalling factors on the right. (C) Model of Crassulaceae subsidiary cell (SC) development; *KlaxMUTEs* establish meri-stemoid2 identity and drive SC divisions by inducing an asymmetric division program and delaying guard cell (GC) differentiation. Stages are indicated and cell identities color-coded; stomatal lineage ground cells (SLGCs) are in lilac, SCs in yellow, guard mother cells (GMCs) and GCs in green, meristemoid1 in purple, meristemoid2 in orange.

*thaliana* (Fig. 4B, Table S3 and S4). In conclusion, some of the downstream genetic modules activated by *MUTE* seem to have diversified in *K. laxiflora* so that GC commitment is delayed and an ACD program required to form SCs is induced.

## Discussion

Much like in grasses, the two *MUTE* homologs adopted a novel function to form SCs in a eudicot leaf succulent. Considering the substantial evolutionary distance between monocot grasses and eudicot Crassulaceae succulents, and the distinct ontogeny of their perigene and mesogene SCs, respectively, this is a remarkable example of a convergent genetic mechanism. We propose that *KlaxMUTE1* and *KlaxMUTE2* control the transition from meristemoid1, which executes a *SPCH*-like ACD program generating SLGCs, to meristemoid2, which generates SCs instead (Fig. 4C). In contrast, *AtMUTE* terminates asymmetric divisions, commits meristemoids to GMCs and promotes their symmetric division (Han et al., 2022, 2018; Pillitteri et al., 2007). The diverse role of the different MUTE proteins, however, seemed not to be encoded within the proteins themselves, as the ACD promoting versus ACD inhibiting function could not be transferred from *A. thaliana* to *K. laxiflora*, or vice versa. Instead, MUTE proteins activate divergent, species-specific downstream genetic programs to modify developmental programs required to generate GCs in *A. thaliana*, and SCs in *K. laxiflora*. This is of little surprise considering that changes to cis-regulatory elements of putative target genes are likely less deleterious and more flexible than mutating the transcription factor itself, and mutations within cis-elements are observed repeatedly as a strategy generating phenotypic divergence within plants and animals (Marand et al., 2023; Wittkopp and Kalay, 2011). In SC-less *A. thaliana, AtMUTE* finetunes cell cycle and GMC commitment by inducing expression of a GMC-specific CycD5;1 and the GC differentiation factor *AtFAMA* to guarantee a single GMC division and GC differentiation (Han et al., 2018; Ohashi-Ito and Bergmann, 2006).

In addition, *AtMUTE*-dependent expression of a cyclin-dependent kinase (CDK) inhibitor (*SIAMESE-RELATED4*) prevents CYCD3;1 and CYCD3;2 from interacting with CDKs, thus preventing ACDs (Han et al., 2022). In SC-forming *K. laxiflora* on the other hand, *KlaxMUTE1* induced specific *CycD3* genes to sustain SC-forming ACDs and downregulated transcription factors like *KlaxFAMA2* and *KlaxFLP* that are associated with GC differentiation and division inhibition in *A. thaliana* (Lai et al., 2005; Ohashi-Ito and Bergmann, 2006).

It remains unclear how meristemoid2 divisions are ended and GMCs specified. The absence of mature GCs in *klaxmute1;klaxmute2* could be due to a role of *KlaxMUTEs* in GMC establishment or a mere consequence of not establishing meristemoid2 identity. Despite the apparent redundancy of *KlaxMUTE1* and *KlaxMUTE2*, their overexpression phenotypes differed slightly. *KlaxMUTE1*^*OE*^ showed lower ploidy and an increase in stomatal density, while stomatal density was slightly decreased in *KlaxMUTE2*^*OE*^, and the effect on ploidy was weaker. Taken together, with the strong induction of *KlaxMUTE2* expression in *KlaxMUTE1*^*OE*^, and a slightly weaker and delayed expression of *KlaxMUTE2* translational reporters in stage 3, our results suggest sequential roles for these proteins. *KlaxMUTE1* might act earlier than *KlaxMUTE2*, which might have a role in the meristemoid2-to-GMC transition.

Also, it is unclear how SCs differentiate in *K. laxiflora*. Potentially, SC differentiation is induced directly, post-mitotically, as a daughter cell of a meristemoid2 division. Alternatively, *KlaxMUTEs*’ presence in SCs might be required during the GMC stage (stage 6-7), and/or the GMC itself might be involved in the establishment of SC identity through cellular signalling processes. Indeed, expression of both *KlaxMUTEs* increased in stage 6-7 SCs, which could be due to reinforced cell-autonomous translation, or additional cell-to-cell mobility, as is known for grasses (Raissig et al., 2017; Wang et al., 2019). *KlaxMUTEs’* translational reporter signal in both nucleus and cytoplasm of GMCs was consistent with those of other mobile transcription factors like BdMUTE (Raissig et al., 2017), or AtSHR (Nakajima et al., 2001).

Finally, staining for K^+^ within open and closed stomata revealed that the SCs of succulent leaves of *K. laxiflora* are indeed functionally relevant, and likely act as stomatal helper cells. In grasses, SCs were shown to act as osmolyte source and sink (Raschke and Fellows, 1971), to accommodate GC movement biomechanically by supporting inverse turgor pressurization (Durney et al., 2023; Franks and Farquhar, 2007). Furthermore, loss of SCs had a severe negative impact on opening and closing speed when they were genetically ablated in the model grass *B. distachyon mute* mutant (Raissig et al., 2017; Spiegelhalder et al., 2024). Unlike in grasses, where the perigene SCs stem from a distinct cell lineage to the one that generates GCs, the *klaxmute1;klaxmute2* double mutant aborted both SC and GC formation as they are consecutively formed by the same cell lineage. To show unequivocally the physiological role of SCs using genetic ablations, even more sophisticated transgene approaches like cell-type-specific amiRNAs or well-timed rescue of GMC formation, will be required. We speculate, however, that SCs might be pressurized when GCs are closed to minimize stomatal conductance efficiently and prevent daytime transpiration. Indeed, the substantial larger total volume of *K. laxiflora* SCs compared to GCs and the fact that SCs are rarely completely devoid of K^+^, even when forced open, indicate a more prominent role of SCs during stomatal closing.

In conclusion, our work shows that, much like in monocots, MUTE can be flexibly deployed to facilitate the development of SCs in CAM succulents in the eudicot family Crassulaceae. Importantly, our work lays the foundation to manipulate experimentally, and dissect mechanistically, developmental aspects of the water-use efficient CAM leaf anatomy and the inverted daily cycle of stomatal opening and closing that supports CAM. Succulence and (inducible) CAM metabolism are largely ignored, yet highly promising traits to engineer water conservation strategies in agricultural systems.

## Materials & Methods

### Plant material & growth conditions

*Kalanchoë laxiflora* accession Oxford Botanical Garden (OBG) was used as wild type throughout this study. To grow seedlings on plates, seeds were surface-sterilized using 70% (v/v) ethanol [Fisher Scientific] for 5 min, washed three times with sterile water, resuspended in 0.1% agarose [BioConcept], and plated on ½ Murashige and Skoog (MS) plates (per liter: 2.2 g Murashige and Skoog medium including Gamborg’s B5 vitamins [Duchefa: M0231], 30 g sucrose [Fisher Scientific: BP220], 8 g Phytoagar [Duchefa: P1003] with pH adjusted to 5.8 using 1 M NaOH). Plates were grown in a Percival plant growth chamber using diel cycles of 16-h-light/ 8-h-dark (45 - 50 μmol PAR m^−2^ s^−1^, light period temperature = 22°C, dark period temperature = 18°C).

For soil-grown plants, either plants with 4-5 leaf pairs (LPs, with LP1 being the first visible, youngest leaf pair) and 1cm long roots were transplanted from plates to soil, or seeds were directly dispersed in a pot (8 cm diameter) with seedling substrate [Klasmann-Deilman GmbH, Germany, Article # 455433]. After around 2-weeks growth under 12-h-light/ 12-h-dark cycles (300 μmol PAR m^−2^ s^−1^; light period temperature = 24°C, dark period temperature = 18°C), single seedlings were transplanted to bigger pots (12 cm diameter) containing the following soil; 5 parts “Landerde” [Ricoter, Switzerland], 4 parts peat and 1 part white sand (hereafter referred as “*Kalanchoë* soil”) and moved to a greenhouse with 16-h-light/ 8-h-dark cycles (300 μmol PAR m^−2^ s^−1^; light period temperature = 24°C, dark period temperature = 18°C).

*Arabidopsis thaliana* Col-0 plants for floral dip transformation were grown in a greenhouse with a 16-h-light/ 8-h-dark cycle (130-150 μmol PAR m^−2^ s^−1^; light period temperature = 22°C, dark period temperature = 18°C). Twenty to thirty *Arabidopsis* seedlings were grown in 6 cm pots containing seedling substrate [Klasmann-Deilmann GmbH, Germany] and used for transformation (see below). To grow and select T0 seedlings on plates, seeds were sterilized three times using 70% (v/v) ethanol, poured onto sterile filter paper, dried and evenly spread on ½ MS plates (per liter: 2.15 g Murashige and Skoog medium [Duchefa: M0221], 30 g sucrose [Fisher Scientific: BP220], 8g Phytoagar [Duchefa: P1003] with pH adjusted to 5.8 using 1M NaOH) containing 50 mg L^-1^ Kanamycin [CarlRoth: T832].

### Vegetative propagation of *Kalanchoë laxiflora*

*K. laxiflora* can be grown through stem cuttings or plantlets that emerge on the margins of leaves that were detached from the mother plant.

Stem cuttings were cut between leaf pair 4 (LP4) and LP5 and planted into Jiffy pots containing *Kalanchoë* soil. After approximately 2-weeks of root growth in Jiffy pots in a glasshouse under 16-h-light/ 8-h-dark cycles (300 μmol PAR m^−2^ s^−1^; light period temperature = 24°C, dark period temperature = 18°C), Jiffy pots were transferred to bigger pots (12 cm diameter) containing *Kalanchoë* soil, and growth was continued in the same conditions.

For plantlet growth, single leaves were detached from the mother plant and placed individually in pots (8 cm diameter) containing *Kalanchoë* soil. Once the plantlets formed around 6 LPs, they were transferred individually to Jiffy pots containing *Kalanchoë* soil. Once they reached a stem length of around 10 cm, they were transferred to big pots (12 cm diameter) containing *Kalanchoë* soil.

### Flower induction in *Kalanchoë laxiflora*

For flower induction, the main stem of *K. laxiflora* must be at least 10 cm long (around 5-6 LPs) and grown in longday conditions (i.e. 16-h-light/ 8-h-dark) for at least 2 weeks. Plants from a glasshouse grown under 16-h-light/ 8-h-dark cycles (300 μmol PAR m^−2^ s^−1^; light period temperature = 24°C, dark period temperature = 18°C) were transferred to a Percival plant growth chamber under short-day 8-h-light/ 16-h-dark cycles (130-150 μmol PAR m^−2^ s^−1^; light period temperature = 22°C, dark period temperature = 18°C). Shortday conditions induced flowering after ∼3 weeks, however, inflorescence formation was only visible after 4-5 weeks of short-day growth. Once flowering was induced, plants were moved to a glasshouse under 16-h-light/ 8-h-dark cycles (300 μmol PAR m^−2^ s^−1^; light period temperature = 24°C, dark period temperature = 18°C) to expedite flower development.

### Crossing *Kalanchoë laxiflora*

Around 5-6 weeks after inflorescence formation, flowers were mature enough to initiate the crossing process. Flower development in *K. laxiflora* is not synchronized, but different stages of flowers can be found within one inflorescence, resulting in a crossing window of approximately 2-weeks. Flower buds that revealed 5- to 10 mm of petals were carefully opened with forceps and all eight anthers were carefully removed without injuring the four stigmas. Anthers should look purple and not have dehisced mature pollen already to avoid self-pollination during emasculation. Petals were closed carefully with forceps after emasculation and emasculated flowers were labelled using tape. Forceps should be cleaned with 70% (v/v) ethanol between each emasculated flower to avoid cross-pollination. Two days after emasculation, flowers were carefully opened using forceps, and mature stigmas were pollinated with mature pollen of the desired genotype. Anthers with mature pollen were obtained from open flowers with white anthers that released pollen.

To produce *klaxmute1;klaxmute2* double mutants four independent transgenic lines of *klaxmute1* and two independent transgenic lines of *klaxmute2* were used for crossing. *klaxmute1* stigmas were pollinated with *klaxmute2* pollen, and vice versa. For crossing the *KlaxMUTEs* translational reporter lines with the plasma membrane marker line, stigmas of four individuals from three independent lines of *KlaxMUTE1:m-Citrine-KlaxMUTE1* and one individual of *KlaxMUTE2:mCi-trine-KlaxMUTE2* were pollinated with *35S:mCherry-AtPIP1;4* pollen.

### Potassium staining of open and closed stomata

Wild-type *K. laxiflora* OBG plants were grown to the 10–12 LP stage. LP6 was collected, the abaxial side of the epidermis was peeled and float-incubated overnight in an opening or closing buffer (50 mM KCl [Sigma-Aldrich] and 10 mM MES-KOH [Sigma-Aldrich]) with 5 μM fusicoccin [ChemCruz] in the dark or 50 μM abscisic acid (ABA [ChemCruz]) in the light, respectively.

Potassium (K^+^) precipitation was conducted based on (Raschke and Fellows, 1971). In short, upon fusicoccin and ABA treatment, leaf peels were placed in 50 mM Ca(NO_3_)_2_ [Fisher Scientific] in water for 90 s and then in a solution with 0.1 mM Ca(NO_3_)_2_ in water for 30 s. The epidermal peels were then washed with water for 2 min and transferred to a solution of sodium cobaltnitrite (Na_3_Co(NO_2_)_6_; 26.67% (w/v) of Co(NO_3_)_2_ [Sigma-Aldrich] and 46.67% (w/v) of NaNO_2_ [Sigma-Aldrich] in a 13% (v/v) solution of acetic acid [Fisher Scientific]) in water for 30 min. The samples were washed three times in deionized water for 2 min each and immersed in a solution of 5% (v/v) ammonium sulfide [ChemCruz] to fix the samples. After a final wash in deionized water for 30 s, samples were mounted on slides for visualization and imaged using a Leica DM2000 LED light microscope with a Leica DMC6200 camera.

To quantify in which stomatal cell K^+^ precipitated, three biological replicates with three technical replicates each were analyzed. Each stomatal complex was assigned to one of five categories depending on K^+^ precipitation location, the stomatal complexes were counted fully anonymized by four people, classified, and the percentage for each classification was calculated. A one-way ANOVA was performed followed by a 5% Tukey’s HSD test (alpha = 0.05) to compare between classes and within treatments and pairwise t-tests to compare within classes but between treatments.

### Molecular Cloning

#### CRISPR/Cas9 constructs

Guide RNA sequences with high on-target activity score and (almost) no off-targets were selected in Geneious (https://www.geneious.com/) targeting *KlaxMUTE1* (Kl-Gene012921, priXC24 and priXC25) and *KlaxMUTE2* (Kl-Gene023418, priXC18 and priXC19). To generate the CRISPR/Cas9 constructs, we used an adapted pFASTRK vector from VIB-UGent’s collection (vector ID: 12_01). The binary vector was digested with BsaI [New England Biolabs] and dephosphorylated with Antarctic Phosphatase [BioConcept] and the single-stranded primer guides with appropriate overhangs were annealed, phosphorylated with T4 kinase [New England Biolabs] and ligated into the linearized backbone.

The guide against *KlaxMUTE1* was intentionally designed to potentially also target *KlaxMUTE2*, but only *klaxmute1* single mutant lines were retrieved. For genotyping, leaf genomic DNA was extracted using a modified CTAB protocol (Raissig et al., 2016). The *klaxmute1* mutation was amplified using priXC61 and priYBG4, the *klaxmute2* mutation was amplified using priXC122 and priXC123, and amplicons were Sanger sequenced. The same primers and approach was used to confirm homozygous mutations in *klaxmute1;klaxmute2* double mutant seedlings.

All primer sequences can be found in Table S1.

#### Reporter & overexpression constructs

All reporter- and overexpression constructs were cloned using the GreenGate system (Lampropoulos et al., 2013). *KlaxMUTEs* promoter and coding sequence were amplified from *K. laxiflora* OBG wild-type genomic DNA extracted using a modified CTAB protocol (Raissig et al., 2016). For creating the entry module *pGGA_KlaxMUTE1pro* and *pGGA_ KlaxMUTE2pro*, priXC6 and priXC7 were used to amplify 1313 bp upstream of *KlaxMUTE1* and priXC59 and priXC11 were used to amplify 2109 bp upstream of *KlaxMUTE2*. The amplicons were digested by BsaI [New England Biolabs] and ligated into the pGGA000 backbone (BsaI digested and dephosphorylated by Antarctic Phosphatase [BioConcept]). To make the entry modules *pGGC_KlaxMUTE1* (including the STOP codon), primers priXC4 and priXC5 were used to amplify the *KlaxMUTE1* genomic sequence. The fragment was digested by BsaI and ligated into the BsaI digested and dephosphorylated backbone pGGC000 by T4 DNA ligase [New England Biolabs]. To create the entry module *pGGC_ KlaxMUTE2* (including the STOP codon), two separate fragments were amplified using primers priXC8/priXC12 and priXC13/priXC9 to mutate the BsaI site in the second intron. Both fragments were digested and ligated into the BsaI digested and dephosphorylated backbone pGGC000. All cloned pGGA and pGGC entry vectors were test digested and whole insert Sanger sequenced.

To generate the transcriptional reporter *KlaxMUTE1p:mCitrine-eGFP*^*nls*^: pGGA_KlaxMUTE1pro, pGGB_ mCitrine_IPK_Ala-linker, pGGC_eGFP_NLS (pGGC012), pGGD_Dummy (pGGD002), pGGE_rbscT (pGGE001) and pGGF_NOSpro-KanR (pGGF007) were assembled into the destination vector pGGZ004 by GreenGate cloning. To build the transcriptional reporter *KlaxMUTE2p:mCitrine-eG-FP*^*nls*^: pGGA_KlaxMUTE2pro, pGGB_mCitrine_IPK_Ala-linker, pGGC_eGFP_NLS (pGGC012), pGGD_Dummy (pGGD002), pGGE_rbscT(pGGE001) and pGGF_NOS-pro-KanR (pGGF007) were assembled into the destination vector pGGZ004 by GreenGate cloning. To generate the translational reporter *KlaxMUTE1p:mCitrine-KlaxMUTE1:* pGGA_KlaxMUTE1pro, pGGB_mCitrine_IPK_Ala-linker, pGGC-KlaxMUTE1, pGGD-Dummy (pGGD002), pGGE_ rbscT (pGGE001) and pGGF_NOSpro-KanR (pGGF007) were assembled into the destination vector pGGZ004 by GreenGate cloning. To build the translational reporter *KlaxMUTE2p:mCitrine-KlaxMUTE2:* pGGA_KlaxMUTE2 pro, pGGB_mCitrine_IPK_Ala-linker, pGGC_KlaxMUTE2, pGGD-Dummy (pGGD002), pGGE_rbscT (pGGE001) and pGGF_NOSpro-KanR (pGGF007) were assembled into the destination vector pGGZ004 by GreenGate cloning. To build the overexpression construct *35S:mCitrine-KlaxMUTE1*: pG-GA_35Sp (pGGA004), pGGB_mCitrine_IPK_Ala-linker, pGGC_KlaxMUTE1, pGGD-Dummy (pGGD002), pGGE_ rbscT (pGGE001) and pGGF_NOSpro-KanR (pGGF007) were assembled into the destination vector pGGZ004 by GreenGate cloning. To produce the overexpression construct *35S:mCitrine-KlaxMUTE2*: pGGA_35Sp (pGGA004), pGGB_mCitrine_IPK_Ala-linker, pGGC_KlaxMUTE2, pGGD-Dummy (pGGD002), pGGE_rbscT (pGGE001) and pGGF_NOSpro-KanR (pGGF007) were assembled into the destination vector pGGZ004 by GreenGate cloning.

All assembled vectors were test digested and whole-plasmid Sanger sequenced or the overhang sites were sequenced. The entry modules pGGA000, pGGC000, pGGA004, pGGC012, pGGD002, pGGE001, pGGF007 were previously described in (Lampropoulos et al., 2013). pGGZ004 was described in (Lupanga et al., 2020). To generate pGGB_mCi-trine_IPK_Ala-linker, the mCitrine including the Ala-linker was amplified from the *BdFAMAp:mCitrine-BdFAMA* construct (McKown et al., 2023) using primers priMR454 and priMR455 and generated as described above for promoter and CDS entry clones.

To build the overexpression construct *35S:mCitrine-AtMUTE(CDS)*: first, primers priTN63 and priTN94 were used to amplify the *AtMUTE* coding sequence from cDNA of *A. thaliana* Col-0 plants, which was synthesized with the PrimeScript RT Master Mix (Perfect Real Time [Takara]) from RNA extracted with the RNeasy Plant Mini Kit [Qiagen]. The entry module pGGC_AtMUTE(CDS) was produced as described above. The binary vector was generated by assembling pGGA_35Sp (pGGA004), pGGB_mCitrine_ IPK_Ala-linker, pGGC_AtMUTE(CDS), pGGD-Dummy (pGGD002), pGGE_rbscT (pGGE001) and pGGF_NOS-pro-KanR (pGGF007) into the destination vector pGGZ004 using GreenGate cloning.

To clone the plasma membrane marker *35S:mCherry-AtPIP1;4*: pGGA_35Sp (pGGA004), pGGB_mCherry-GSL, pGGC_AtPIP1;4, pGGD-Dummy (pGGD002), pGGE_rb-scT (pGGE001) and pGGF_NOSpro-KanR (pGGF007) were assembled into the destination vector pGGZ003 by GreenGate cloning. The pGGB_mCherry-GSL and pGGC_ AtPIP1;4 were generously provided by Karin Schumacher and Alexis Maizel (COS, Heidelberg University, Germany), respectively. pGGZ003 was previously described in (Lampropoulos et al., 2013).

All primer sequences can be found in Table S1.

#### Tissue-culture-based transformation of *Kalanchoë laxiflora* accession OBG diploid

The stable transformation of *K. laxiflora* OBG diploid followed the protocols described in (Dever et al., 2015) and (Boxall et al., 2020), but adapted as follows.

For tissue-culture-based transformation, around 10 *K. laxiflora* seedlings per plate were prepared as described above. Around 5 plates per construct were used for transformation.

To prepare the bacteria cell culture, 4 mL of LB plus the required antibiotics (here: 50 mg L^-1^ Rifampicin [Sigma-Aldrich], 25 mg L^-1^ Gentamicin [Sigma-Aldrich] and 50 mg L^-1^ Spectinomycin [Sigma-Aldrich]) were inoculated with a single colony of the *Agrobacterium tumefaciens* strain GV3101 carrying the construct of interest and incubated overnight at 28ºC with rotational shaking at 180 rpm. The overnight culture was pelleted at 3000 g for 10 min at room temperature in a 50 ml Falcon tube. The pellet was gently resuspended in around 20 mL MS30 media (per liter: 4.41 g Murashige and Skoog medium including Gamborg’s B5 vitamins [Duchefa: M0231], 30 g Sucrose [Fisher Scientific: BP220]) and the OD600 adjusted to 0.1. Finally, 40 mL of the MS30 media and bacteria solution was used for transformation. Acetosyringone [Sigma-Aldrich] was added to a final concentration of 100 μM before the tube was wrapped with aluminum foil and incubated with gentle shaking for at least 2-h at room temperature to induce GV3101 virulence genes that mobilize the transfer-DNA (T-DNA).

Meanwhile, the four biggest leaves of the seedlings were harvested into a sterile petri dish using sterile scissors in a sterile laminar flow tissue culture cabinet. Each leaf was cut in two to three pieces using a sterile scalpel with a minimum leaf piece width of around 5 mm. Once all leaf pieces were cut, they were placed into the Falcon tube containing the *A. tumefaciens* cell suspension with the construct of interest. The leaf pieces were incubated in the *A. tumefaciens* cell suspension for approximately 1-h at room temperature with gentle shaking. Afterwards, most of the *A. tumefaciens* solution was removed and the leaf pieces were placed onto fresh Callus Induction Medium (CIM) plus Acetosyringone [Sigma-Aldrich] plates (per liter: 4.41 g Murashige and Skoog medium including Gamborg’s B5 vitamins [Duchefa: M0231], 30 g Sucrose [Fisher Scientific: BP220], 8 g Phytoagar [Duchefa: P1003], 1 mg L^-1^ TDZ [Duchefa: T0916] and 0.2 mg L^-1^ IAA [Duchefa: I0901] with the pH adjusted to 5.7 using 1M NaOH) plus 100 µM Acetosyringone [Sigma-Aldrich] but with no antibiotics for initial co-cultivation. Leaf pieces were placed on CIM + Acetosyringone plates, wrapped in aluminium foil, and grown for 48-h at 22ºC light period temperaturem (16-h), 18ºC dark period temperature (8-h). After this step, plant tissues were grown in a Percival plant growth cabinet under 16-h-light/ 8-h-dark cycles (45-50 μmol PAR m^−2^ s^−1^; light period temperature = 22°C, dark period temperature = 18°C). Leaf pieces were then transferred to fresh CIM plates with added antibiotics (per liter: 4.41 g Murashige and Skoog medium including Gamborg’s B5 vitamins [Duchefa: M0231], 30 g Sucrose [Fisher Scientific: BP220], 8 g Phytoagar [Duchefa: P1003], 1 mg L^-1^ TDZ [Duchefa:T0916], 0.2 mg L^-1^ IAA [Duchefa: I0901], 100 mg/ L^-1^ Kanamycin [Carl-Roth: T832] and 300 mg L^-1^ Timentin [Duchefa: T0190] with the pH adjusted to 5.3 using 1M NaOH) every two weeks until callus formation (approx. 8- to 12-weeks). At each of the first two subculturing iterations, small plantlets that formed on the leaf margins of the leaf pieces were excised and discarded. Once callus was visible on single leaf pieces, those pieces were moved to fresh Shoot Induction Medium (SIM) plates (per liter: 4.41g Murashige and Skoog medium including Gamborg’s B5 vitamins [Duchefa: M0231], 30 g Sucrose [Fisher Scientific: BP220], 8 g Phytoagar [Duchefa: P1003], 1 mg L^-1^ BAP [Duchefa: B0904], 0.2 mg L^-1^ IAA [Duchefa: I0901], 100 mg L^-1^ Kanamycin [CarlRoth: T832] and 300 mg L^-1^ Timentin [Duchefa:T0190] with the pH adjusted to 5.1 using 1M NaOH). Callus pieces were subcultured every two weeks to fresh SIM plates for shoot development (approx. 8-weeks). Shoots that developed from the same callus were numbered and treated as clones. Callus pieces containing several shoots or single shoots were cut into smaller pieces or excised below the stem, respectively, and placed on Root Induction Medium (RIM) plates (per liter: 4.41g Murashige and Skoog medium including Gamborg’s B5 vitamins [Duchefa: M0231], 30 g Sucrose [Fisher Scientific: BP220], 8 g Phytoagar [Duchefa: P1003], 50 mg L^-1^ Kanamycin [CarlRoth: T832] and 300 mg L^-1^ Timentin [Duchefa:T0190] with the pH adjusted to 5.2 using 1M NaOH). Shoots were subcultured every two weeks until roots of approximately 1 cm had developed. Regenerated plants with 3-4 LPs and a developed root system were transferred to Jiffy pots containing *Kalanchoë* soil and grown in a greenhouse under 16-h-light/ 8-h-dark cycles (300 μmol PAR m^−2^ s^−1^; light period temperature = 24°C, dark period temperature = 18°C).

### Plant transformation of *Arabidopsis thaliana*

*Arabidopsis thaliana* Col-0 wild-type plants were transformed with the *35S:mCitrine-KlaxMUTE1, 35S:mCitrine-KlaxMUTE2* and *35S:mCitrine-AtMUTE* constructs, respectively, using the *A. tumefaciens* strain GV3101-based floral dip transformation protocol (Clough and Bent, 1998).

### Brightfield and confocal microscopy

To image the mature epidermis samples, the abaxial side of LP5 (around 3 cm long) was carefully peeled and moved to a microscopy slide. The epidermal peel was stained with Toluidine Blue O solution (0.2 g Toluidine Blue O [ChemCruz] in 40 mL Acetate Buffer (0.01 g pectolyase [Duchefa: P8004] in 84.7 mL 1M acetic acid [Fisher Scientific] and 10 mL distilled water)) for a few seconds and washed with water at least three times until the water on the slide was transparent. The epidermal peel was mounted on a microscopy slide in 50% glycerol [Fisher Scientific]. The samples were imaged on a Leica DM2000LED [Leica Microsystems] using brightfield settings.

To quantify the overexpression lines compared to wild type, three fields of view of three independent individuals per genotype were used. Wild-type-like mature stomatal complexes, mature stomatal complexes with more than three SCs and the total number of cells per field of view were counted in each image using Fiji (Schindelin et al., 2012). To calculate stomatal density and total cell density, number of mature stomatal complexes and total number of cells were divided by the size of the field of view 0.198 mm^2^, respectively. The percentage of mature stomatal complexes with more than three SCs was calculated by dividing the number of mature stomatal complexes with more than three SCs by the total number of mature stomata. For plotting and statistical analysis, the values from three fields of view per individual were averaged.

For *klaxmute1;klaxmute2* phenotyping, cotyledons of plate-grown *klaxmute1;klaxmute2* and phenotypic wild-type-like seedlings were carefully collected 14 days after sowing and the cell membrane of the leaf epidermis was stained with 0.01mM FM4-64 [Sigma-Aldrich]. After staining for 5 min, cotyledons were rinsed with water and mounted on a microscopy slide with water. All samples were imaged with a Leica Stellaris 5 confocal microscope [Leica Microsystems]. FM4-64 signal was excited using 549 nm (∼10-20% intensity) of a white light LED laser running at 85%. Single slice, mid-plane images were taken with a 63X glycerol objective at 1024×1024 pixels and a line average of three.

To image the *35Sp:mCitrine-KlaxMUTE1, 35Sp:mCi-trine-KlaxMUTE2* and *35Sp:mCitrine-AtMUTE* overexpression lines in *A*.*thaliana*, plate-grown cotyledons of seedlings were collected around 11 days after sowing and stained with FM4-64 as described above. To image the *35Sp:mCitrine-KlaxMUTE1, 35Sp:mCitrine-KlaxMUTE2* and *35Sp:mCitrine-AtMUTE* overexpression lines in *K*.*laxiflora*, the 2^nd^ leaf pair (around 0.5 cm) was collected and stained with FM4-64 as described above. To image overexpression lines in *A*.*thaliana* and *K*.*laxiflora*, single-slice, mid-plane images were taken with a 63X glycerol objective at 1024×1024 pixels and a line average of three. For FM4-64 excitation, 549 nm (∼10-20% intensity) of a white light LED laser running at 85% was used. For mCitrine excitation, 515 nm (∼10% intensity) of a white light LED laser running at 85% was used.

To image transcriptional and translational reporter lines, 0.5- to 2 cm leaves (mostly 2^nd^ or 3^rd^ leaf pair) were used. To achieve better FM4-64 staining and confocal imaging quality, cuticular wax was removed by gently rubbing the leaf with some water containing hand sanitizer or soap. The leaf was cut next to the mid vein with a scalpel and only part of the leaf was used for imaging. For transcriptional reporter lines, leaves were stained with FM4-64 as described above. The translational reporter lines were crossed with the plasma membrane reporter so no staining was needed. To avoid sample damaging, leaf samples were surrounded by four lines of vaseline before mounting in water. The cover slip was carefully placed onto the vaseline having the leaf sit in a water-filled chamber. Single-slice, mid-plane images were taken with a 63X glycerol objective at 1024×1024 pixels and a line average of three. For FM4-64 or mCherry excitation, 549 nm (∼10-20% intensity) of a white light LED laser running at 85% was used. For mCitrine excitation, 515 nm (∼10% intensity) of a white light LED laser running at 85% was used.

### Manual time-lapse imaging

The *K. laxiflora* plasma membrane marker line (*35Sp:m-Cherry-GSL-AtPIP1;4*) was used for manual time-lapse imaging. The plants were between 10 to 15 cm tall for the stem to be flexible enough for mounting but not too long to risk stem breaking. The pots of the plants were wrapped in cling foil to avoid soil contamination of the microscope. The leaf pair to be imaged was selected (ideally 0.5 - 1 cm) and the second leaf of the pair and any shoot growing above it were removed with clean scissors, as well as other leaves inferring with mounting. The leaf to be imaged was taped to a small plastic petri dish using micropore tape and part of the edge of the plate was cut out for the stem to lay flat. A spot of Lanolin mixed with charcoal was placed on the leaf with a pipet tip, preferably in a relatively flat area between leaf veins. This mark was used as a reference to find the imaged area again. The whole plant was placed on a styrofoam box next to the stage of the Leica Stellaris 5 confocal microscope and the petri dish with the prepared leaf was placed below the objective. A 10X water dip-in lens was used and the stage was adjusted to have the lanolin mark right under the centre of the objective before placing some water between the sample and the objective. In the brightfield channel the leaf epidermis was focused and a notable spot at the edge of the lanolin mark was selected. Then the channel was switched to fluorescence, and starting at the lanolin landmark, Z-stacks covering the entire depth of the epidermis were taken for four overlapping tiles (with an estimated 10% overlap). For mCherry excitation, 549 nm (∼8% intensity) of a white light LED laser running at 85% was used and the images were taken in a 704×704 pixel format, Z-stack step size 0,41 µm and a line average of 1. The Z-stacks were converted to 3D-objects in Fiji (Schindelin et al., 2012) and oriented to a ‘flat’ position. The correctly oriented images were saved in .png format and stitched manually using Inkscape (version 1.1 (c68e22387, 2021-05-23)).

### Ploidy analysis

Four adult leaves (around 4 cm) were used per sample and two samples were prepared for each genotype (wild type, *KlaxMUTE1*^*OE*^, *KlaxMUTE2*^*OE*^). LoBind reaction tubes [Eppendorf] were used for all steps. The abaxial side of the leaf was peeled off with forceps and placed in 250 µL pre-cooled CyStain UV Precise P Nuclei Extraction Buffer [Sysmex]. The peeled epidermis of all four leaves per genotype were pooled in one tube and incubated for 1 min on ice. A 50 µm filter [Sysmex] was pre-treated with 250 µL 1% BSA [Sigma-Aldrich] in Phosphate-buffered saline (PBS, per liter: 8 g NaCl [Fisher Scientific], 0.2 g KCl [Sigma-Aldrich], 1.44 g Na_2_HPO_4_ [ChemCruz], 0.24 g KH_2_PO_4_ [ChemCruz] with pH adjusted to 7.4). Next, the extracted nuclei were collected by pipetting up and down three times and filtered through the filter. Then, the filter was washed with 500 µL CyStain UV Precise P Staining Buffer (containing DAPI [Sysmex]). The flow-through was collected in the same tube as the extracted nuclei. The samples were kept on ice and gently mixed before Flow Cytometry analysis with the BD FACS LSR II system. The data was analysed with the BD FACSDiva software (version 8.0.1). The counts for both samples per genotype were combined.

### Isolation of high molecular weight total genomic DNA from *K. laxiflora* OBG diploid for genome sequencing

For the isolation of high molecular weight (HMW) total genomic DNA for whole genome sequencing, a single *K. laxiflora* OBG diploid plant was grown and sampled. The single plant sampled was grown from a batch of seed that had passed through three rounds of self-pollination and single seed descent in order to increase the homozygosity of the genome. The plant was grown to maturity (∼30 cm tall) over 6-months in a research glasshouse under 16-h-light/ 8-h-dark cycles with a minimum temperature in the light period of 20°C. Prior to leaf sampling for HMW total genomic DNA isolation, the plant was dark-adapted in constant darkness at 15°C for 3-days to ensure all leaf starch had been turned over. Leaf pairs (LP) 1 and 2 (where LP1 are the first LP that are large enough to handle, either side of the shoot apical meristem) were then collected from multiple shoot tips of the dark adapted plant, snap frozen in liquid nitrogen and stored at -80°C until use for genomic DNA isolation.

The frozen, dark-adapted LP1 and LP2 samples were pooled together, ground to fine powder in liquid nitrogen using a pestle and mortar, and then 100 mg of powdered leaf tissue was used for each total genomic DNA isolation using the Cytiva Nucleon PhytoPure 0.1 g kit [Cytiva RPN8510] according to the manufacturer’s protocol. The quality and quantity of the DNA was analyzed initially using a Nanodrop spectrophotometer and the average size distribution was checked using agarose gel electrophoresis to ensure the DNA was intact and of a high average molecular weight, greater than ∼30 Kb. The HMW total genomic DNA sample was submitted to the University of Liverpool Centre for Genomic Research (CGR) for QC analysis and library preparation for PacBio HiFi long-read sequencing.

### PacBio library preparation and long-read sequencing

The HMW genomic DNA from *K. laxiflora* OBG diploid was analyzed by the CGR using a Thermofisher Qubit fluorometer for accurate quantification of the dsDNA, and an Agilent Fragment Analyzer for confirmation that the DNA was of HMW, averaging above ∼30 Kb.

DNA was then used for PacBio DNA library preparation, and the library was sequenced on the Sequel system using the PacBio SMRT flow cell-8; the library was sequenced on 3 cells to generate > 100-fold genome coverage. The PacBio sequencing chemistry 3.0 was used and the resulting raw reads were used for genome assembly and annotation, as described below. The PacBio raw reads (BAM files) are available from the NCBI Short Read Archive (SRA) under the Bioproject ID PRJEB83678.

### Genome assembly

The raw sequence reads from the 3 PacBio flow cells were filtered to retain only the reads that were 1 kb or longer, which resulted in 1,744,005 reads that spanned a total of 26.3 Gbp with a median read length of 10,324 bp. The filtered reads were assembled using Canu v1.8 (Koren et al., 2017), yielding an estimated genome size of 260 Mbp, which was consistent with the prediction from flow cytometry analysis of 274 Mb (Flow Cytometry Services, Netherlands). The contigs resulting from the Canu *de novo* assembly were polished using the PacBio raw reads and the packages pbalign (https://github.com/PacificBiosciences/pbalign) and Arrow (Chin et al., 2013), with 5 iterations. The final assembly metrics are summarised in Table S5.

Prior to annotation the assembled genome was assessed for completeness using BUSCO v2.0 (Simão et al., 2015) using database species “arabidopsis” and database lineage “embryophyte_odb9”. The results from the BUSCO analysis are summarised in Table S6.

### Genome Annotation

Gene annotations across the 78 contigs from the Canu assembly were compiled using MAKER (Holt and Yandell, 2011), which uses both multiple sources of published evidence and *ab initio* prediction to generate gene models.

### Repeat modelling and repeat masking of the genome

Repeat sequences including LTRs and MITEs were indentified and modelled in the assembled genome using RepeatModeler v1.0.11 (Smit et al., n.d.), MiteFinder v1.0.006 (Hu et al., 2017), LTRharvest (Ellinghaus et al., 2008) and LTRdigest (Steinbiss et al., 2009) in the GenomeTools package v1.5.11 (Gremme et al., 2013). In addition to the genome sequence, various public databases were utilised in the process, namely: the UniProt UniRef protein database, a eukaryotic tRNA database and a transposase database.

### Generation of AUGUSTUS *ab initio* gene models

The *ab initio* gene predictor AUGUSTUS (Stanke et al., 2006) is an integral component of the MAKER pipeline. Here, AUGUSTUS v3.2.3 was used. An initial run of MAKER was performed supported by the following evidence: Uniprot/swissprot proteins, *K. laxiflora* (tetraploid; accession number 1982-6028) 309 v1.1 proteins (from the public database Phytozome), *K. fedtschenkoi* 382 v1.1 proteins (from Phytozome; (Yang et al., 2017)), in-house unpublished proteins from *K. fedtschenkoi* accession “Kew-Glasgow-Liverpool” from Liverpool CGR project LIMS17639, and *K. laxiflora* (tetraploid; accession number 1982-6028) 309 v1.1 transcripts (from Phytozome). MAKER was instructed to predict gene models directly based on transcript and protein evidence mappings (est2genome=1, protein2genome=1). This generated predictions that were generally unrefined, but those that were most accurate were used to train AUGUSTUS iteratively. The resulting species-specific Hidden Markov Model (HMM) was used in the final MAKER run.

### Annotation with MAKER

MAKER v2.31.9 was used to generate a raw set of gene models with the following evidence and parameters. As part of the MAKER run, the software was instructed to repeat-mask the genome using both generic repeat models and the generated genome-specific repeats.

Evidence: Uniprot/swissprot proteins, *K. laxiflora* (tetraploid; accession number 1984-6028) 309 v1.1 proteins (from the public database Phytozome), *K. fedtschenkoi* 382 v1.1 proteins (from Phytozome; (Yang et al., 2017), in-house unpublished proteins from *K. fedtschenkoi* accession “Kew-Glasgow-Liverpool” from Liverpool CGR project LIMS17639, and *K. laxiflora* (tetraploid; accession number 1984-6028) 309 v1.1 transcripts (from Phytozome), plus the AUGUSTUS HMM generated as described above. The “Run Arguments” set were as follows: augustus_species=Kl, keep_preds=1. The AUGUSTUS models were used to predict genes and all predictions were retained (i.e. whether predictions were supported by provided evidence or not) as these were filtered downstream. This initial annotation with MAKER yielded 30,141 raw gene models.

### Basic functional annotation

Raw protein sequences generated from the raw gene models produced by MAKER were annotated with basic functional information from Unitprot/swissprot protein alignments that were generated using BLASTP.

### Filtering gene models

Raw gene models were filtered to generate a final set of gene models. First, protein sequences derived from the MAKER gene models were annotated using InterProScan v5.31-70.0 (Jones et al., 2014), which detected Pfam domains and Gene Ontology (GO) terms. Gene annotations were updated with the InterProScan results. Raw gene models were then filtered and retained if they fulfilled two criteria: (a) they were supported by RNA or protein evidence, or (b) they had no supporting RNA or protein evidence but were predicted to contain a Pfam domain. This filtering step reduced the final set of predicted genes to 27,768.

### Functional annotation

The final set of 27,768 gene models were annotated with the eggNOG-mapper v1.0.1 (Huerta-Cepas et al., 2017) using the eukaryotic functional database of eggNOG. In addition, the final 27,768 predicted genes were used to re-assess genome completeness using BUSCO v2.0 as already described above. Table S7 summarises the comparison between the initial BUSCO analysis of the assembled genome contigs and the final BUSCO analysis of the 27,768 gene models that were retained after complete annotation and filtering.

### Assessment of duplication based on the final, filtered gene set

In addition to BUSCO analysis providing an estimate of genome/ transcriptome completeness and the level of duplication further analyses were carried out to explore the level of duplication in the genome based on the final filtered gene set. The protein sequences predicted from the final set of 27,768 genes were used as input for cd-hit-est to cluster proteins at 99% identity, and the cluster sizes resulting from this were recorded. Clusters of size 2 or greater would suggest duplication. In addition, clusters were classified as being in the same genomic neighbourhood if 25% of genes in the cluster (minimum 2) were within 50 kb of the genes in the cluster. High proportions of proteins in clusters that included 2 or more protein sequences could indicate redundancy in the genome assembly or may be caused by true duplicates within the genome. In addition, clusters with 2 or more genes in the same neighbourhood would indicate high levels of local gene duplication. The clusters recovered from this analysis are summarised in Fig. S8. This analysis demonstrated that protein sequences derived from the final set of filtered gene models displayed a very low level of duplication across the *K. laxiflora* OBG diploid genome.

### RNA sequencing of wild type and *KlaxMUTE1*^*OE*^

For bulk-RNA sequencing, the 7^th^ leaf pair was harvested from four wild-type individuals and three independent *KlaxMUTE1*^*OE*^ lines. The samples were snap-frozen in liquid nitrogen and total RNA was extracted using the RNeasy Plant Mini Kit with optional on-column DNAse digest [Qiagen]. The quantity and quality of the purified total RNA was assessed using a Thermo Fisher Scientific Qubit 4.0 fluorometer with the Qubit RNA BR Assay Kit [Thermo Fisher Scientific, Q10211] and an Advanced Analytical Fragment Analyzer System using a Fragment Analyzer RNA Kit [Agilent, DNF-471], respectively. Sequencing libraries were made with 500 ng input RNA using a Revelo mRNA-Seq for MagicPrep NGS kit B [Tecan, PN 30186622] according to the Revelo mRNA-Seq for MagicPrep NGS User Guide [Tecan publication number MO1535, v1] with 15 cycles of PCR. The resulting cDNA libraries were evaluated using a Thermo Fisher Scientific Qubit 4.0 fluorometer with the Qubit dsDNA HS Assay Kit [Thermo Fisher Scientific, Q32854] and an Agilent Fragment Analyzer [Agilent] with a HS NGS Fragment Kit [Agilent, DNF-474], respectively. Pooled cDNA libraries were sequenced paired-end using an illumina NovaSeq 6000 SP Reagent Kit v1.5 (200 cycles; [illumina, 20040719]) on an illumina NovaSeq 6000 instrument. The quality of the sequencing run was assessed using illumina Sequencing Analysis Viewer (illumina version 2.4.7) and all base call files were demultiplexed and converted into FASTQ files using illumina bcl2fastq conversion software v2.20. The quality control assessments, generation of libraries and sequencing was conducted by the Next Generation Sequencing Platform, University of Bern. Approximately 23 - 33 million reads were generated per sample (Table S3). RNA-seq data have been deposited in GEO under accession code GSE.

The reads were mapped against the *K. laxiflora* OBG genome using HISAT2 and counted using htseq-count on usegalaxy.org. Differentially expressed genes were identified using DESeq2 (Love et al., 2014). Genes with a padj < 0.1 and a log2FoldChange>1 or log2FoldChange<-1 were considered differentially expressed. To compare it to induced *AtMUTE* overexpression (i.e. *iMUTE*), we reanalyzed the dataset from (Han et al., 2018; GEO accession number GSE107018) using the above-described pipeline. We identified the *K. laxiflora* OBG homologs of *A. thaliana* stomatal development genes (GO0010374 and own curated list) using local blast against the *K. laxiflora* OBG proteome in Geneious (https://www.geneious.com/).

## Supporting information

Table S1. Primer sequences

Table S2. Count data

Table S3. RNA-seq analysis KlaxMUTE1 overexpression

Table S4. RNA-seq re-analysis iMUTE

---

**Table S5.**
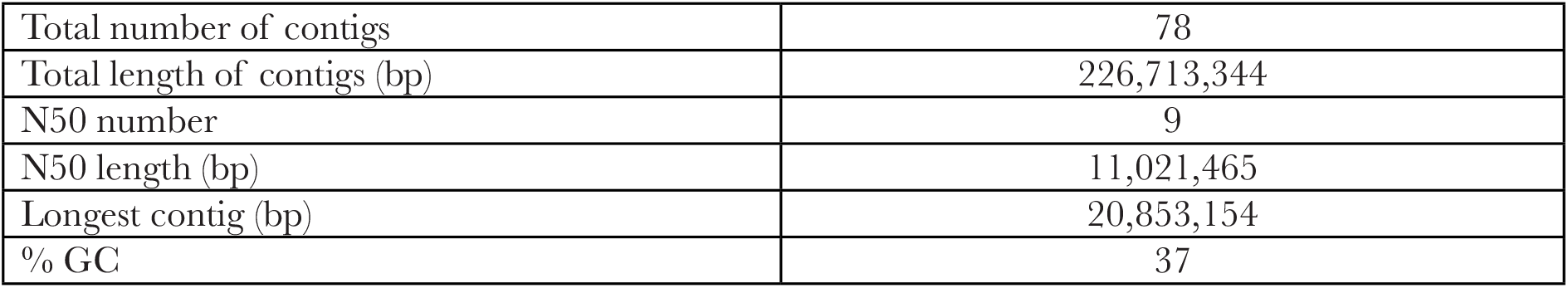
Summary of final assembly metrics for the *de novo* PacBio-only genome assembly of *K. laxiflora* OBG diploid.

|  |  |
| --- | --- |
| Total number of contigs | 78 |
| Total length of contigs (bp) | 226,713,344 |
| N50 number | 9 |
| N50 length (bp) | 11,021,465 |
| Longest contig (bp) | 20,853,154 |
| % GC | 37 |

**Table S6.** Genome completeness for the *K. laxiflora* OBG diploid genome assessed using BUSCO v2.0.

|  | Number of BUSCOs | % of BUSCOs |
| --- | --- | --- |
| Complete BUSCOs | 1298 | 90.1% |
| .....as single copy | 1185 | 82.3% |
| .....as duplicated copies | 113 | 7.8% |
| Fragmented BUSCOs | 35 | 2.4% |
| Missing BUSCOs | 107 | 7.4% |

**Table S7.** Genome completeness comparison of BUSCO analysis of the initial assembled genome contigs in comparison to the final filtered set of predicted gene models.

|  | % of BUSCOs in assembled genome contigs | % of BUSCOs in final, filtered gene set |
| --- | --- | --- |
| Complete BUSCOs | 90.1% | 89.7% |
| .....as single copy | 82.3% | 81.1% |
| .....as duplicated copies | 7.8% | 8.6% |
| Fragmented BUSCOs | 2.4% | 3.3% |
| Missing BUSCOs | 7.4% | 7.0% |

## Acknowledgements

We are grateful to the research gardeners Jasmin Sekulovski, Christopher Ball, Sarah Dolder and Michael Schillbach for their invaluable support in developing horticultural protocols for *K. laxiflora*. We thank Xiaohan Yang for sending us *K. laxiflora* FTBG cuttings (not used here) and Karin Schumacher and Alexis Maizel for providing cloning vectors. For sequencing support, we acknowledge the Next Generation Sequencing Platform at the University of Bern (RNA-seq) and the Centre for Genomic Research (CGR) at U. Liverpool (Genome-seq) with Margaret Hughes (CGR) in particular for preparing and sequencing the PacBio library. This work is supported by a Chinese Scholarship Council fellowship (to XC), a Swiss Government Excellence Fellowship (to AANGF), the DFG Emmy Noether fellowship RA3117/1-1 (to MTR) and the University of Bern. The genome was supported by the U.S. Department of Energy Office of Science, Genomic Science Program (grant DE-SC0008834) and in part by the Biotechnology and Biological Sciences Research Council (BBSRC grant BB/F009313/1 to J.H.). J.H.P. was supported by a PhD studentship funded by the BBSRC as part of the Newcastle-Liverpool-Durham Doctoral Training Partnership (DTP). The contents of this article are solely the responsibility of the authors and do not necessarily represent the official views of the DOE.

## List of Supplementary Materials

Figure S1 - S8

Table S1 - S4

Movie S1 and S2

### Data availability

RNA-seq data have been deposited in GEO under accession code GSE.

PacBio raw reads (BAM files) are available from the NCBI Short Read Archive (SRA) under the Bioproject ID PR-JEB83678.

The assembled and annotated genome is available on Zenodo with DOI 10.5281/zenodo.14563707.

