## Supplemental Figures S1 - S8 for "MUTE drives asymmetric divisions to form stomatal subsidiary cells in Crassulaceae succulents"

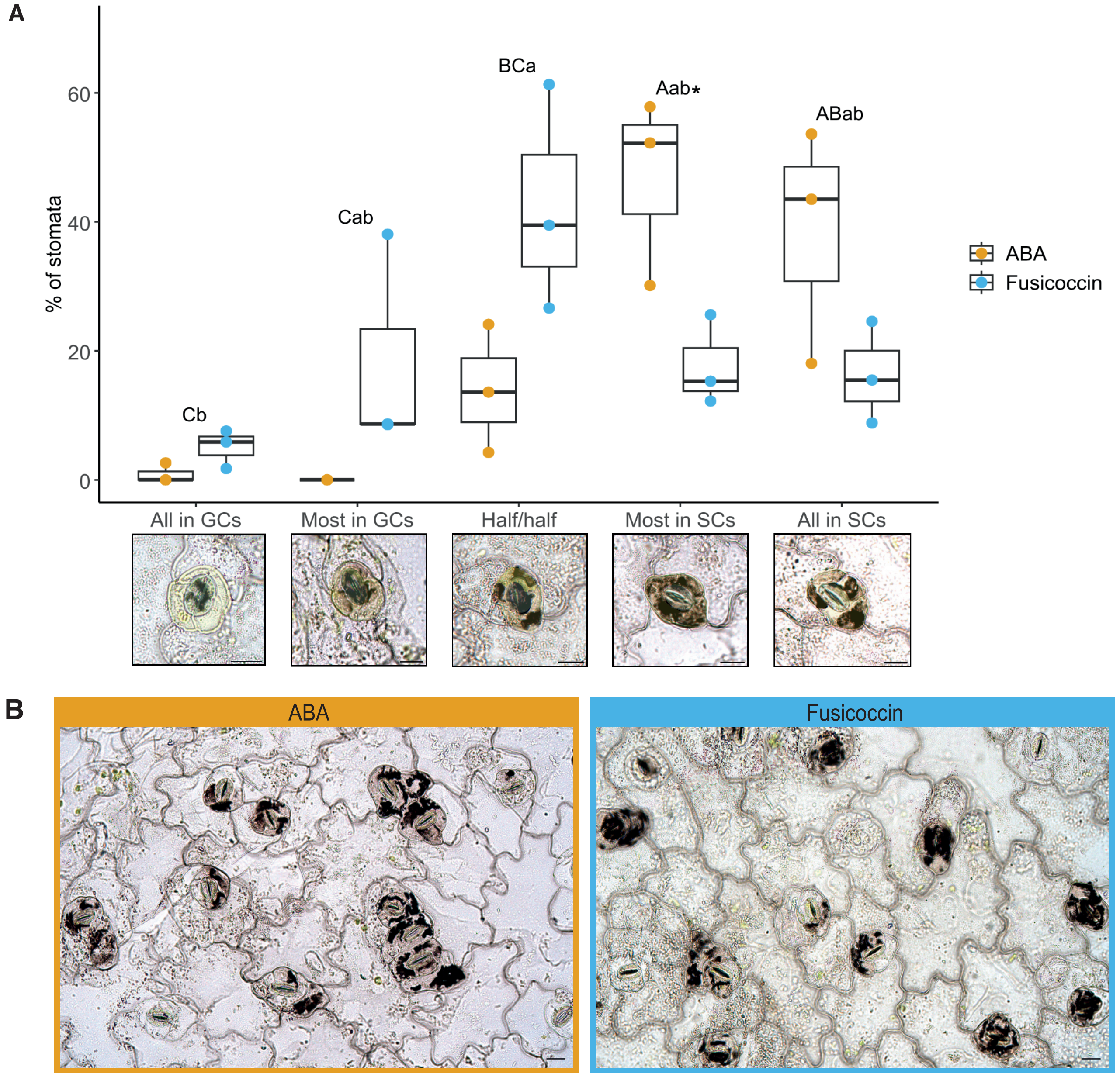


**Figure S1.** Quantification of potassium ($K^+$) levels in open and closed stomata. (A) Statistical analysis of $K^+$ quantification in stomatal complexes of *K. laxiflora* subjected to stomatal opening (Fusicoccin) and closing (ABA) treatment followed by sodium cobaltinitrite treatment for $K^+$ visualization. Differences within a treatment were analysed with a one-way ANOVA followed by Tukey post-hoc test, and capital letters indicate significance groups for ABA and lowercase letters indicate significance groups for Fusicoccin. Pairwise t-test was used to test differences between treatments per class and significance is indicated by an asterisk. Each classification is defined by a representative image of the respective class and the stomata were artificially colored in yellow (subsidiary cells) and green (guard cells); 3 individuals and 84-92 stomata were counted per treatment. (B) Overview of an epidermal leaf peel after ABA treatment (left) and Fusicoccin treatment (right) and $K^+$ staining. Scale bar, 20 $\mu$m.

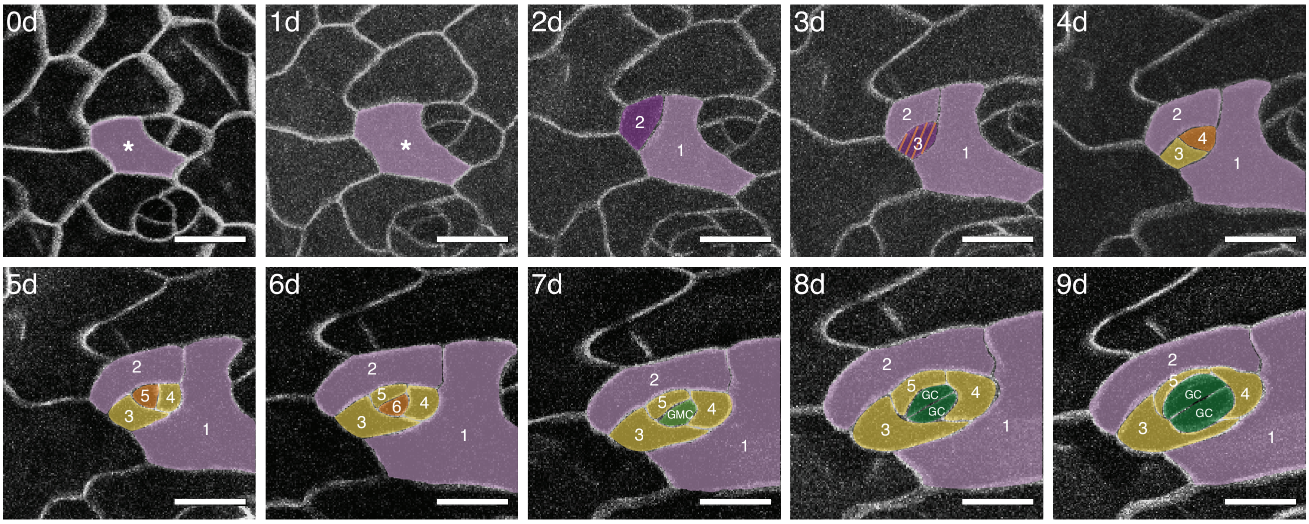


**Figure S2.** Time-lapse imaging of spacing division and consecutive stomatal divisions. 10 day manual time-lapse imaging of plasma membrane marker line *35Sp:mCherry-AtPIP1;4*. Stomata originating from stomatal lineage ground cells (SLGCs) omit the entry division and, consequently, only make one more meristemoid1, SLGC-producing division before transitioning to meristemoid2. Days and cell identities are indicated; SLGCs are in lilac, subsidiary cells (SCs) in yellow, guard mother cells (GMCs) and guard cells (GCs) in green, meristemoid1 in purple, meristemoid2 in orange. Scale bar, 20 $\mu$m.

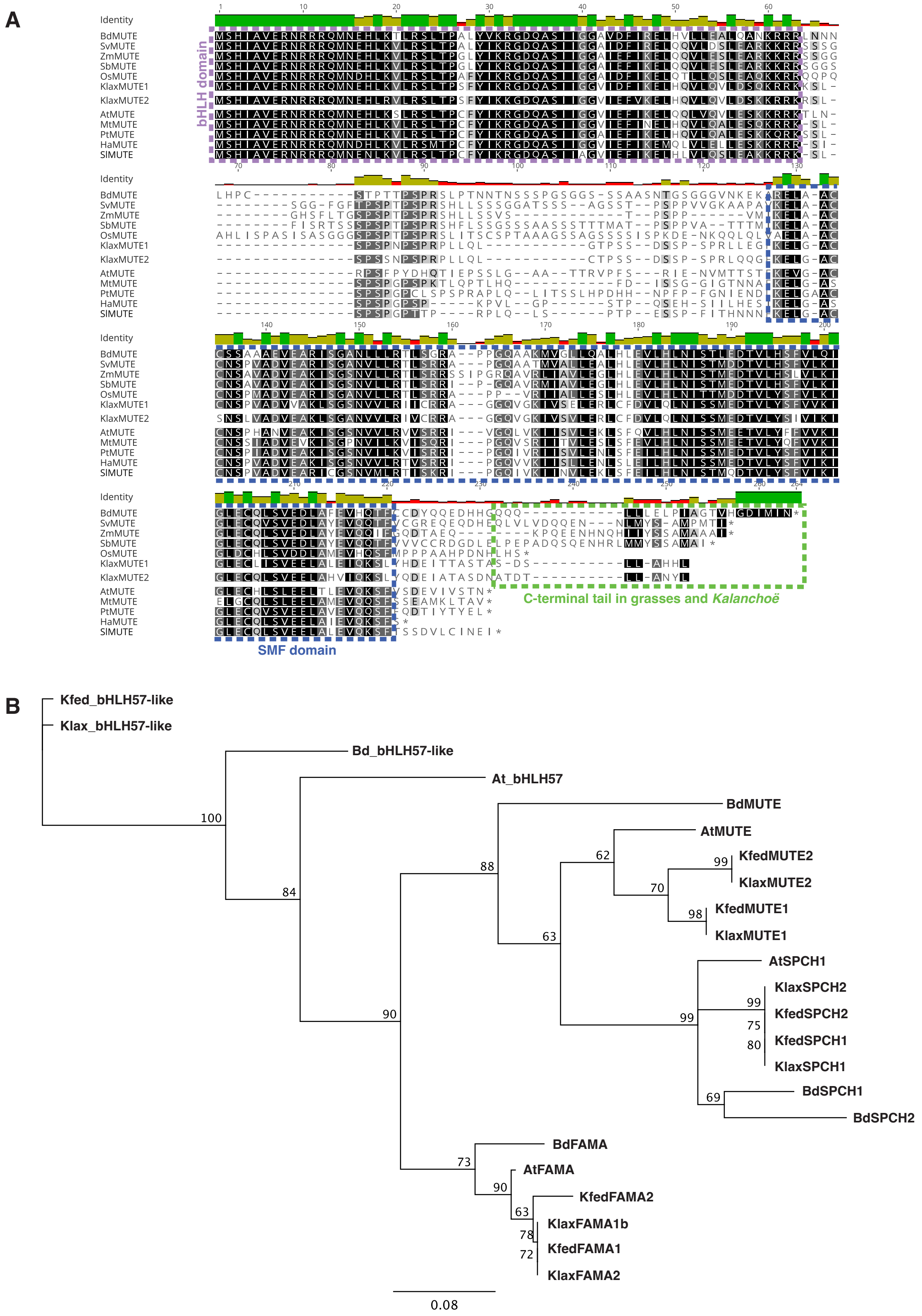


**Figure S3:** MUTE protein alignments and phylogenetic tree of stomatal transcription factors. (A) MUTE protein alignment using Clustal Omega. We used five grass and 6 eudicot species representing the major phylogenetic clades. Grass MUTE protein sequences from *B. distachyon* (Bradi1g18400), *O. sativa* (LOC_Os05g51820), *S. bicolor* (Sobic.009G260200), *Z. mays* (GRMZM2G417164) and *S. viridis* (Sevir.5G150600). Eudicot MUTE protein sequences from *A. thaliana* (At3G06120), *H. annuus* (HanXRQChr12g0362731), *M. truncatula* (Medtr2g022280), *P. trichocarpa* (Potri.008G202900), *S. lycopersicum* (Solyc01g080050) and *K. laxiflora* (KlGene012921 and KlGene023418). (B) Phylogenetic tree of stomatal bHLH transcription factors and bHLH57. Sequences were aligned and the 63 amino acid-long bHLH domain was isolated. A neighbor-joining tree of the isolated bHLH domains with 100 bootstrap reiterations was generated with standard settings in Geneious. No outgroup was specified. Sequences are from *B. distachyon* (BdMUTE = Bradi1g18400, BdSPCH1 = Bradi1g38650.1, BdSPCH2 = Bradi3g09670.1, BdFAMA = Bradi2g22810.1, Bd_bHLH57-like = Bradi1g71990.2), *A. thaliana* (AtMUTE = At3G06120, AtSPCH =AT5G53210.1, AtFAMA = AT3G24140.1, At_bHLH57 = AT2G46810.1), *K. laxiflora* (KlaxMUTE1 = KlGene012921, KlaxMUTE2 = KlGene023418, KlaxSPCH1 = KlGene017537, KlaxSPCH2 = KlGene027982, KlaxFAMA1b = KlGene012763, KlaxFAMA2 = KlOBG012781), *K. fedtschenkoi* (KfedMUTE1 = Kaladp0110s0002.1, KfedMUTE2 = Kaladp0058s0199.1, KfedSPCH1 = Kaladp0058s0411.1, KfedSPCH2 = Kaladp0055s0329.1, KfedFAMA1 = Kaladp0081s0065.1, KfedFAMA2 = Kaladp0081s0067.1, Kfed_bHLH57-like = Kaladp0066s0113.1). KlaxFAMA1a was excluded from the analysis. It does not exist in *K. fedtschenkoi* and is not expressed in developing *K. laxiflora* leaves suggesting it to be a pseudogene or wrongly annotated. The Klax_bHLH57-like gene is from the phytozome *K. laxiflora* FTBG accession (Kalaxd.09G089900.1).

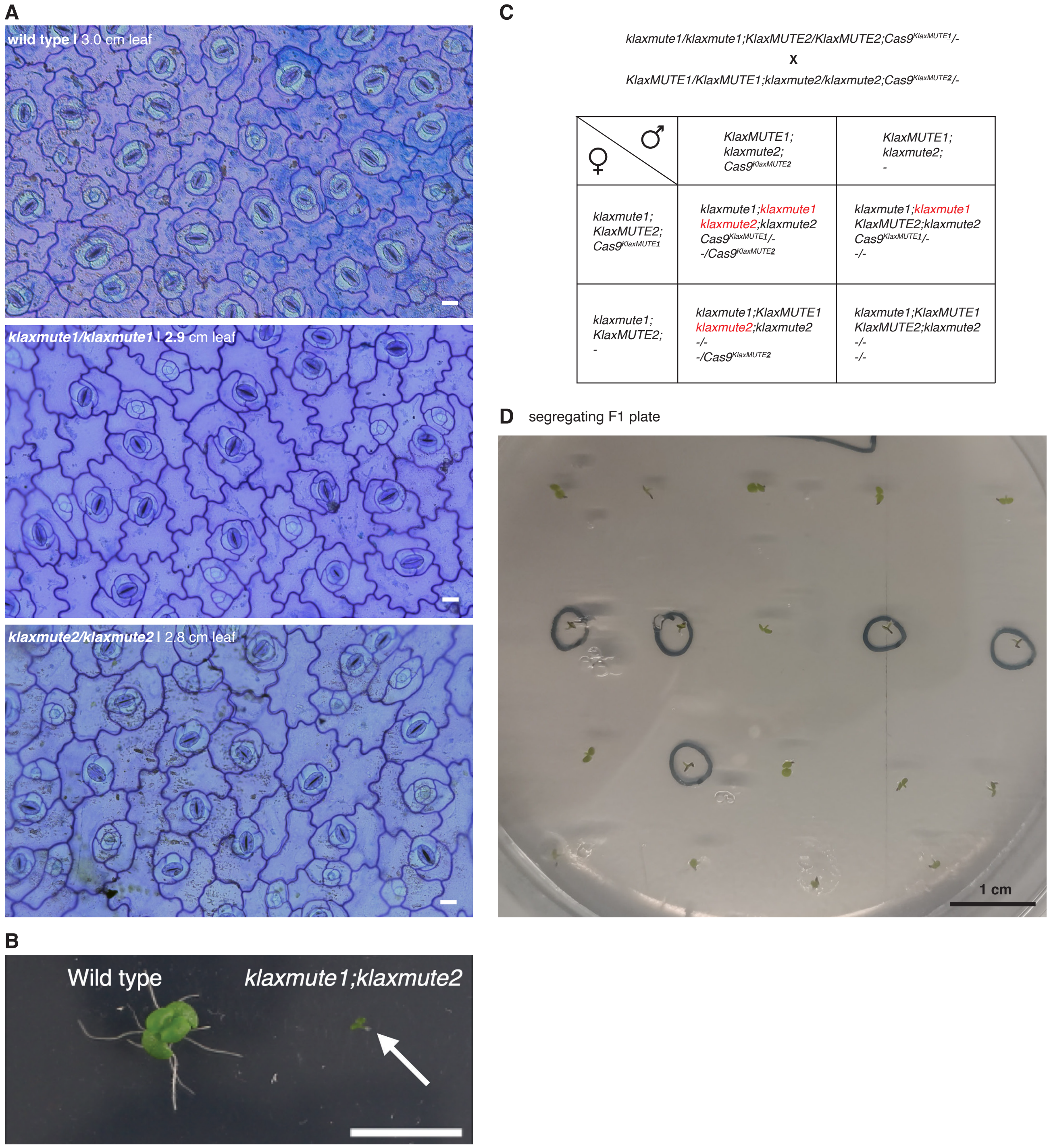


**Figure S4:** *KlaxMUTE1* and *KlaxMUTE2* might have redundant functions in stomatal development. (A) Brightfield images of Toluidine Blue-stained, abaxial epidermal peels of ~3 cm mature leaves from wild-type (top), *klaxmute1* (middle), and *klaxmute2* (bottom) plants. (B) 25d-old seedlings of wild type and *klaxmute1;klaxmute2* double mutants grown on the same ½ MS plate. (C) Punnett square of F1 double mutant screening from crossing *klaxmute1* with *klaxmute2* gene-edited lines that contain an active, hemizygous Cas9 and guideRNA construct. (D) Segregating F1 seedlings on ½ MS plate. The double mutant, arresting seedlings are encircled in black and the wild-type-like seedlings (double heterozygous or homozygous;heterozygous for *KlaxMUTE1* and/or *KlaxMUTE2*) are not encircled. For exact genotypes, see Punnet square in (C). Scale bar, 1cm.

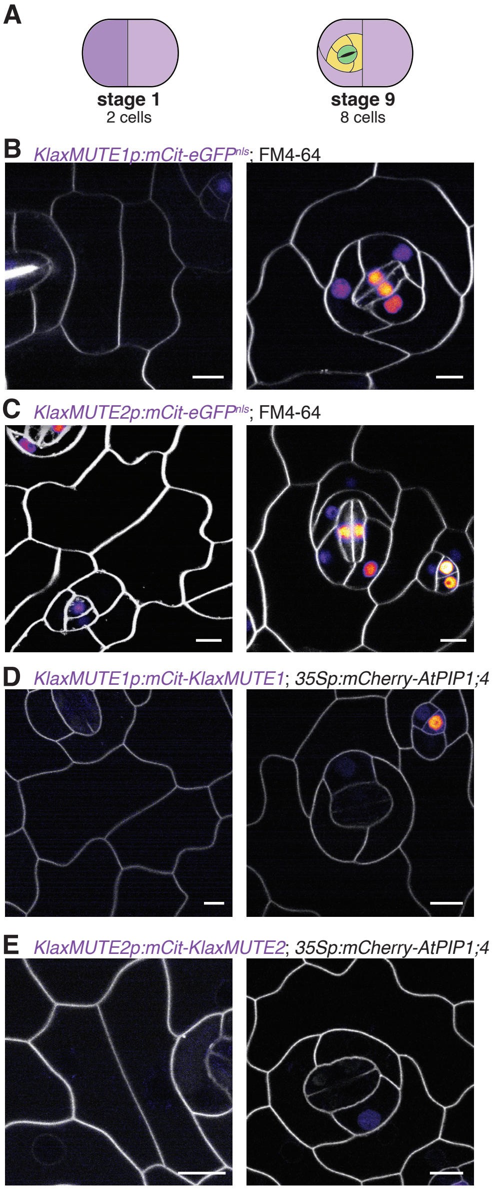


**Figure S5:** Early and mature stages of *KlaxMUTE1* and *KlaxMUTE2* reporter lines. (A) Different developmental stages (stage 1 and stage 9) shown below. (B, C) Confocal microscopy images of transcriptional reporters *KlaxMUTE1p:mCitrine-eGFP*$^{nls}$ (B) and *KlaxMUTE2p:mCitrine-eGFP*$^{nls}$ (C). Cell membrane was stained with FM4-64. (D, E) Confocal microscopy images of translational reporters *KlaxMUTE1p:mCitrine-KlaxMUTE1* (D) and *KlaxMUTE2p:mCitrine-KlaxMUTE2* (E) also expressing the plasma membrane marker *35Sp:mCherry-AtPIP1;4*. Scale bars, 10 $\mu$m.

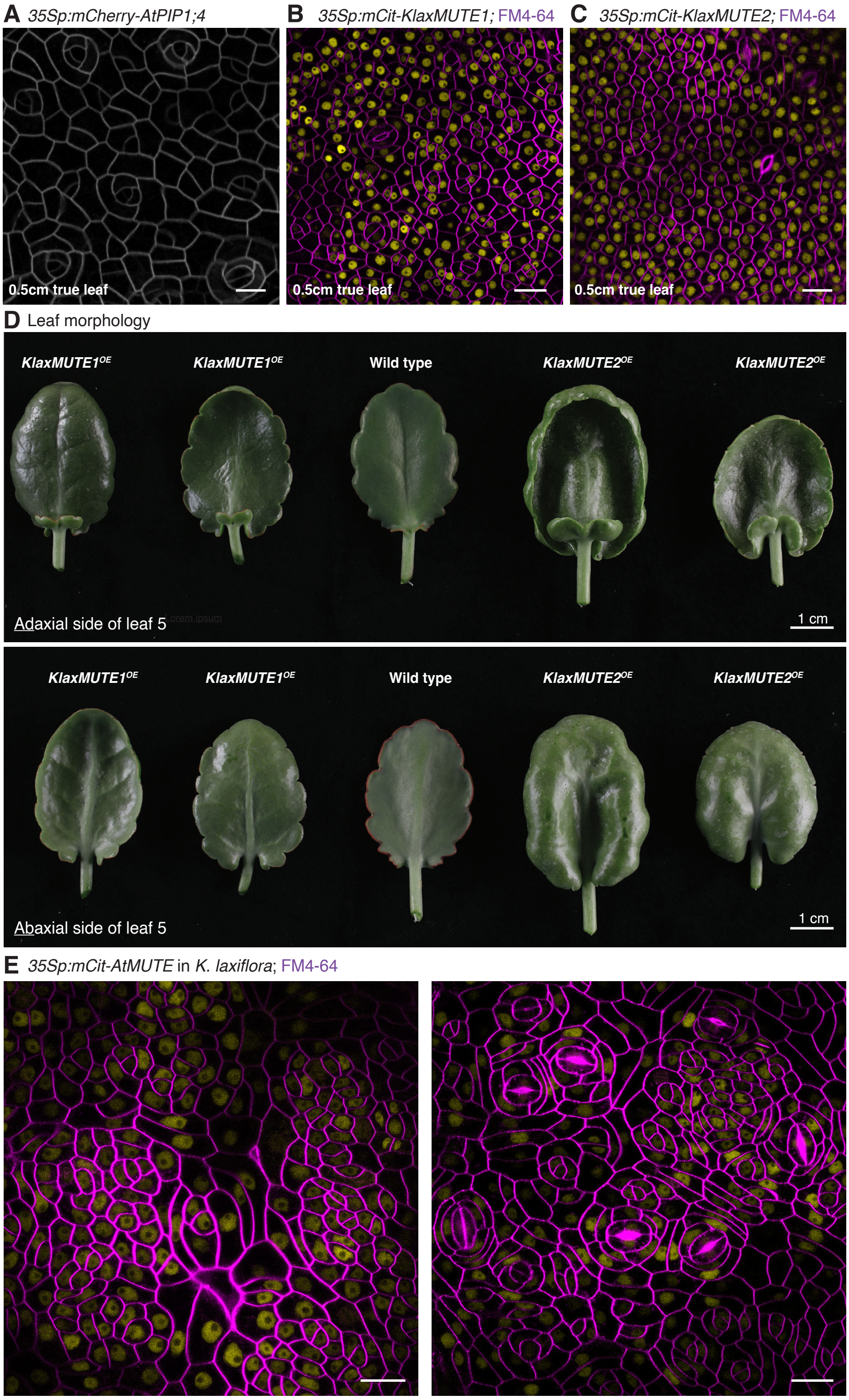


**Figure S6:** Developmental and organ-level phenotype of *KlaxMUTE* overexpression lines in *K. laxiflora*. (A-C) Confocal images of the developing abaxial epidermis of ~0.5 cm leaves are shown from *35Sp:mCherry-AtPIP1;4* (in wild type) (A), *35Sp:mCitrine-Klax-MUTE1* (B), *35Sp:mCitrine-KlaxMUTE2* (C). Cell membrane is stained with FM4-64 in B and C. Scale bar, 20 $\mu$m. (D) *K. laxiflora* leaf morphology of adaxial side (top panel) and abaxial side (bottom panel) of leaf 5 of *35Sp:mCitrine-KlaxMUTE1* (first and second leaf), wild type (middle leaf) and *35Sp:mCitrine-KlaxMUTE2* (fourth and fifth leaf). Scale bar, 1 cm. (E) *35Sp:mCitrine-AtMUTE* expression in *K. laxiflora* leaves induces subsidiary cell-like, asymmetric divisions in the epidermis. Scale bar, 20 $\mu$m.

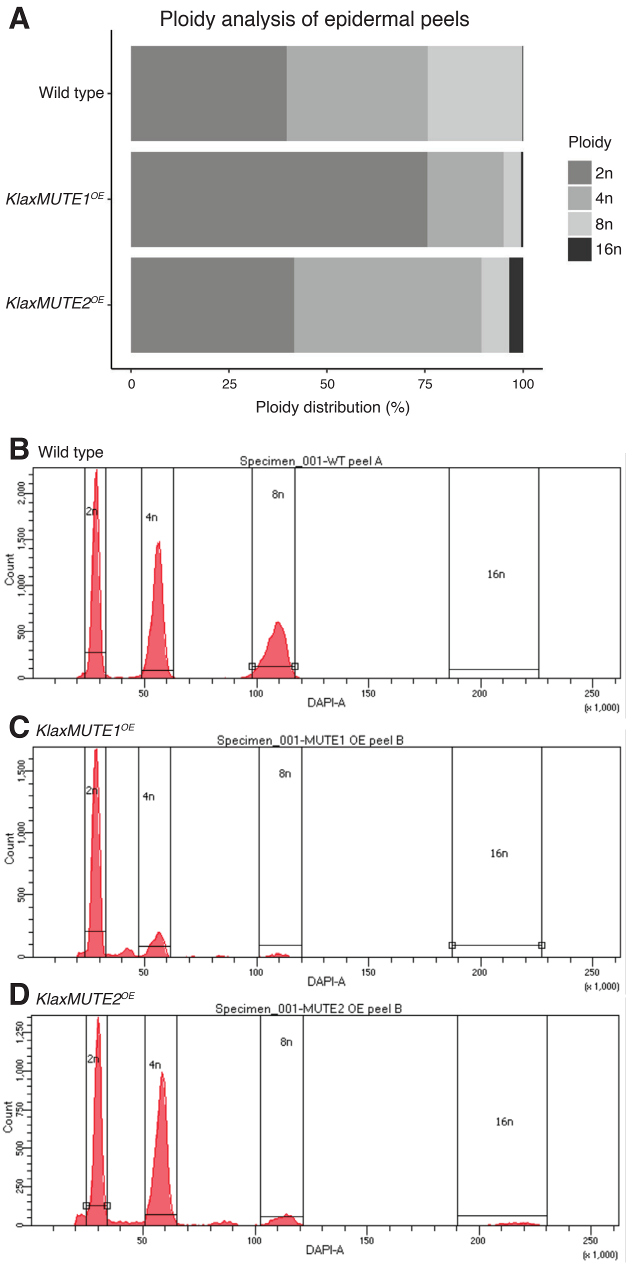


**Figure S7:** Ploidy analysis of peeled leaf epidermis of wild type and *KlaxMUTEs* overexpression lines in *K. laxiflora*. (A) Quantified ploidy levels from flow cytometry runs of isolated and DAPI-stained nuclei from eight adult leaf epidermal peels per genotype. (B-D) Representative individual flow cytometry runs of wild type (B), *35S:mCitrine-KlaxMUTE1* (C), *35S:mCitrine-KlaxMUTE2* (D) are shown and gating is indicated.

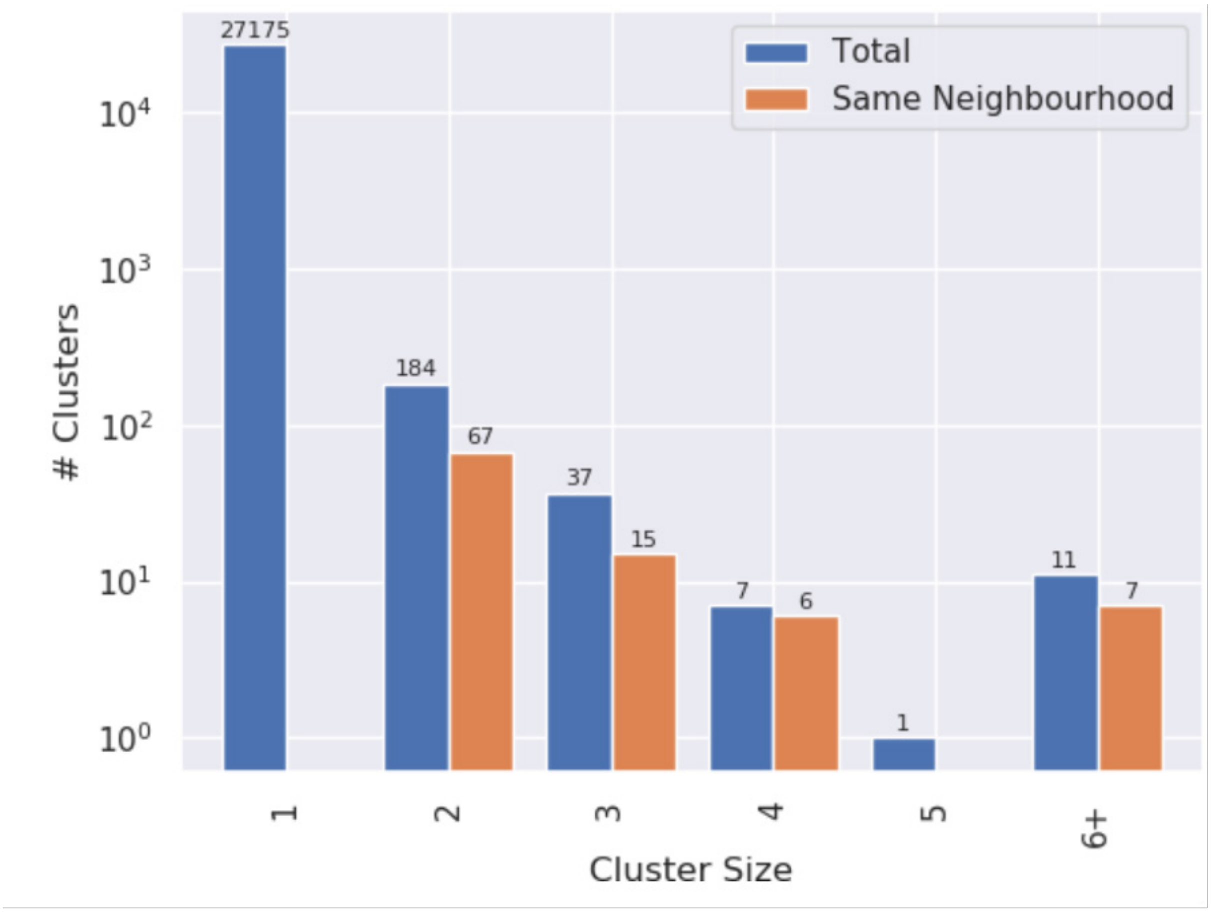


**Figure S8:** Genome-wide analysis for clusters of proteins that were greater than 99% pairwise identity revealed very low levels of gene duplication across the *K. laxiflora* OBG diploid genome assembly and annotation. The number of near identical proteins found to belong to clusters of various sizes (x-axis) were calculated as described in the methods. Blue bars show the number of proteins that were found to belong to clusters of a specific size; where '1' signifies proteins that did not cluster indicative that there was no duplication of that protein. Orange bars show the number of the clusters of each size that were found to be localised within the same genomic region.
