## Supplementary figures and images for "MUTE drives asymmetric divisions to form stomatal subsidiary cells in Crassulaceae succulents"

### Movie S1

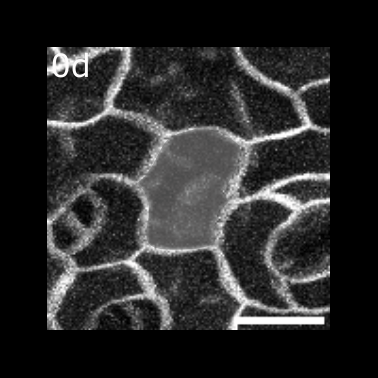

### Movie S2

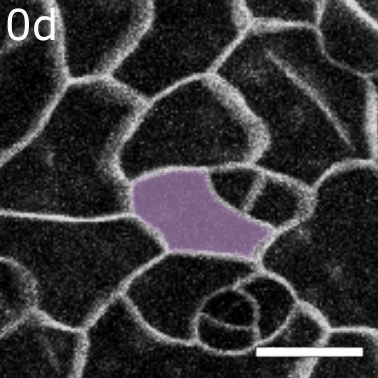
